# Single-cell transcriptomics reveals zone-specific alterations of liver sinusoidal endothelial cells in cirrhosis

**DOI:** 10.1101/2020.03.18.997452

**Authors:** Tingting Su, Yilin Yang, Sanchuan Lai, Jain Jeong, Yirang Jung, Matthew McConnell, Teruo Utsumi, Yasuko Iwakiri

## Abstract

Dysfunction of liver endothelial cells (ECs), particularly sinusoidal endothelial cells (LSECs), is permissive for the progression of liver fibrosis/cirrhosis and responsible for its clinical complications. Here, we have mapped the spatial distribution of heterogeneous liver ECs in normal versus cirrhotic mouse livers and identified zone-specific transcriptomic changes of LSECs associated with liver cirrhosis using single-cell RNA sequencing technology. We identified 6 clusters of liver EC populations including 3 clusters of LSECs, 2 clusters of vascular ECs and 1 cluster of lymphatic ECs. To add finer detail, we mapped the 3 clusters of LSECs to Zones 1 to 3. We found that heterogeneous liver EC identities are conserved even in liver cirrhosis and that Zone 3 LSECs are most susceptible to damage associated with liver cirrhosis, demonstrating increased capillarization and decreased ability to regulate endocytosis. Altogether, this study deepens our knowledge of the pathogenesis of liver cirrhosis at a spatial, cell-specific level, which is indispensable for the development of novel therapeutic strategies to target the most highly dysfunctional liver ECs.

## Introduction

Liver endothelial cells (ECs), including liver sinusoidal endothelial cells (LSECs), vascular ECs and lymphatic ECs, play a central role in liver homeostasis by, among other functions, regulating intrahepatic vascular tone, immune cell function, and quiescence of hepatic stellate cells. Recent development of single-cell RNA sequencing technology has enabled us to identify heterogeneity of these ECs, leading us to link specific EC subpopulations to particular EC functions. Another important factor that can confer different traits to liver ECs is their spatial distributions. The liver consists of repeating anatomical units termed lobules. In each liver lobule, blood flows from the portal vein and hepatic artery toward the central vein, creating gradients of oxygen, nutrients and hormones. In line with these graded microenvironments, key genes in hepatic cells, such as hepatocytes, hepatic stellate cells (HSCs) and ECs, are differentially expressed along the lobule axis, a phenomenon termed zonation^(1–4)^. Therefore, the roles that hepatic cells play in liver physiology and pathophysiology can be zone-specific^(3)^. Thus, characterizing hepatic cells according to their spatial distribution is key to a complete understanding of their physiological functions.

Recognizing the importance of this type of analysis, a recent study by MacParland et al. revealed transcriptomic profiles of heterogeneous hepatic EC populations from healthy human donor livers using scRNA-seq technology and identified three EC populations, including Zone1 LSECs, Zone2/3 LSECs and vascular ECs^(5)^. Another recent study demonstrated the zonation patterns of liver EC genes in mice by paired-cell RNA sequencing, which profiled gene expression of hepatocytes and loosely attached adjacent ECs and determined localization of the ECs in liver lobules based on expression of hepatocyte zonal landmark genes^(1)^. Although special localization was not explored, Ramachandran et al. performed extensive scRNA-seq analyses of all liver non-parenchymal cells, including liver ECs, isolated from human cirrhotic livers in the setting of liver transplantation and determined detailed transcriptomic profiles that were altered in cirrhosis^(6)^.

While these studies have significantly advanced our understanding of heterogeneous EC populations in normal and cirrhotic livers, further characterizations of liver EC populations are still needed to understand important questions related to liver EC biology in both normal and diseased livers. For example, the spatial distribution of liver EC populations has not yet been determined in cirrhosis. Thus, It is not clear whether unique zonal profiles of liver ECs are maintained or lost in cirrhosis. Related to this question, if unique EC populations appear in liver cirrhosis, what are the origins of these cells? How are liver EC transcriptomic profiles altered in liver cirrhosis related to LSEC phenotypes observed in cirrhosis, such as capillarization, EC dysfunction (e.g., dysregulation of vascular tone) and endothelial-to-mesenchymal transition (EndMT)? What are appropriate markers to represent these phenotypic changes in LSECs? Are these phenotypic changes in LSECs zone-specific? How similar or different are these transcriptomic profiles between human and mouse liver ECs in normal and cirrhotic conditions?

To address these questions and others, we performed scRNA-seq analysis of liver ECs isolated from EC-specific green fluorescent protein (GFP) reporter mice, which allowed us to enrich liver EC populations efficiently and exclusively. We first identified heterogeneous liver EC populations. Second, we determined the spatial landscape of these ECs in normal and cirrhotic livers^(1)^. Third, focusing on LSEC populations, we mapped three unique clusters of LSECs that aligned with Zones 1, 2 and 3, determined transcriptomic changes of LSECs in cirrhotic livers in a zone-specific manner, and related these transcriptomic changes to known phenotypic changes of LSECs in cirrhotic livers.

## Results

### Single-cell RNA-seq identified clusters of liver ECs in control and cirrhotic mice

We performed 10x scRNA sequencing analysis on liver EC enriched populations isolated from control and cirrhotic mice. All mice used were EC-specific enhanced green fluorescent protein (eGFP) expressing mice (Cdh5-Cre+, mTmG mice; called EC GFP reporter mice hereafter). Figure 1A illustrates a workflow of cell isolation. GFP-positive and non-apoptotic liver ECs were selected from non-parenchymal cell fractions pooled from 3 mice per group by FACS and were confirmed by a fluorescent image of GFP expression (Figure 1B). Figure 1C illustrates a workflow of data analysis. After excluding low quality cells (expressing fewer than 200 genes or having a mitochondrial genome transcript ratio >0.2) and GFP-negative cells, 3248 cells from control mice and 4076 cells from cirrhotic mice were used for further analysis. Our analysis identified a total of 12 clusters with similar landscapes between control and cirrhotic groups (Figure 1D). Although all the analyzed cells were positive for GFP and VE-cadherin (Cdh5; genes known to be expressed in all ECs) (Figure 1E), some clusters also expressed markers of hepatocytes (Alb, Ttr, Apoa1 and Apoa2), T cells (Nkg7,Trbc1,Trbc2 and Cxcr6), cholangiocytes (Spp1,Krt8, Krt18 and Krt19), macrophages (CD68, C1qa, C1qb and C1qc) and hepatic stellate cells (Colec11, Reln, Dcn and Lrat) (Supplemental Figure 1). Inclusion of other cell types with GFP expression could be due to adherence of ECs to those cells, which may have allowed them to be recognized as single cells during the 10x scRNA seq analysis^(1)^. We excluded these clusters for further analysis and focused only on those clusters with pure EC populations, which included Clusters 1 to 6, corresponding to EC1 to EC5 and lymphatic EC (Figure 1F). The representative marker genes of these clusters are presented with a heatmap (Figure 1G).

**Figure 1.**
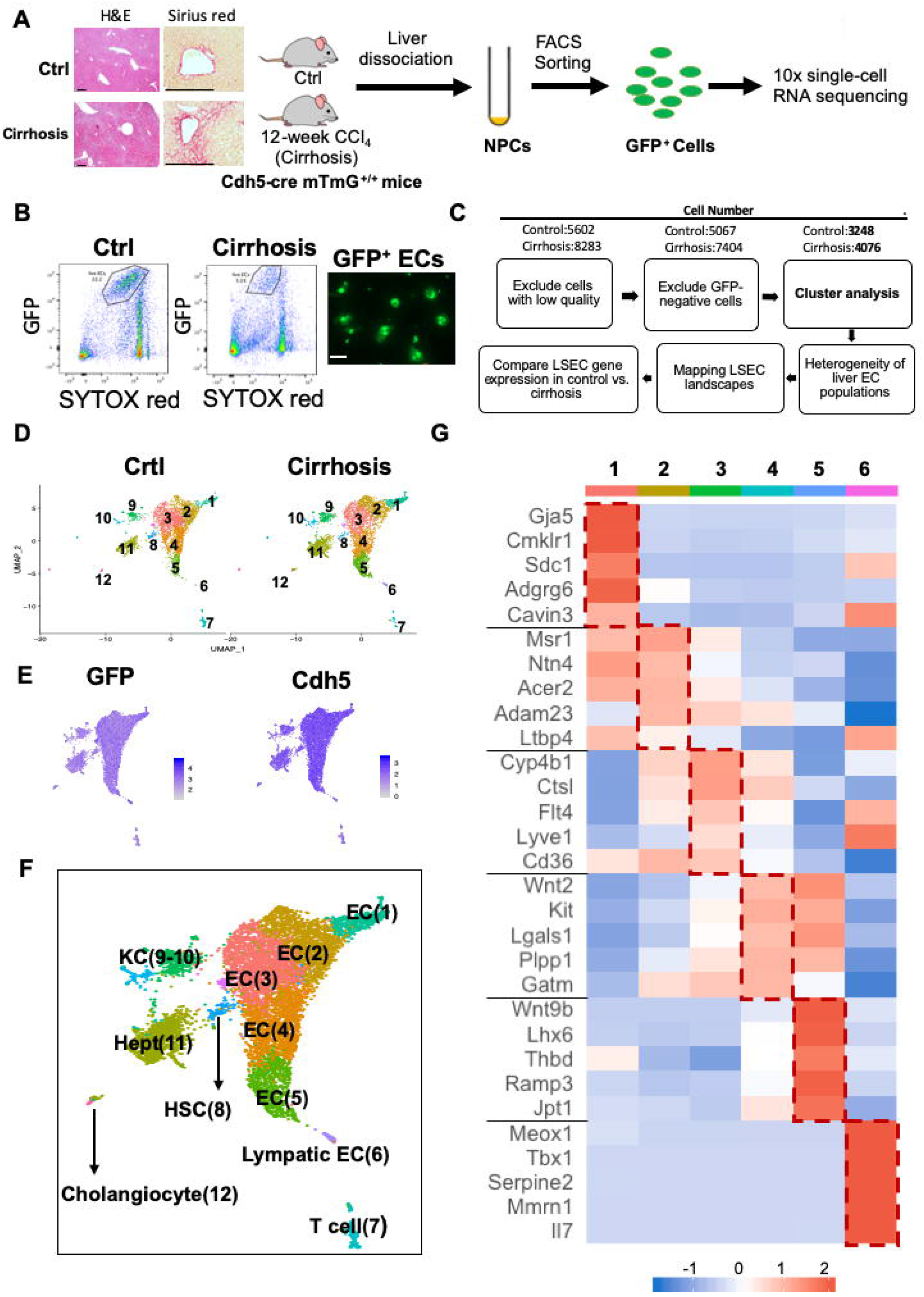
Single-cell RNA-seq revealed a landscape of sorted liver ECs. **A.** Cell isolation workflow using endothelial-GFP reporter mice (tamoxifen inducible, cdh5-cre mTmG^+/+^ mice) subjected to carbon tetrachloride (CCl_4_) inhalation for 12 weeks to generate liver cirrhosis. Age-mached endothelial-GFP reporter mice were used as controls. Representative H&E and Sirius red staining images to show liver injury and fibrotic nodules. Scale bars: 200μm. NPCs: non-parenchymal cells. **B.** Non-parenchymal cells isolated from endothelial-GFP reporter mice were sorted to collect viable GFP-positive cells (i.e., endothelial cells: ECs). SYTOX red staining was used to exclude dead cells (SYTOX red positive cells, left two panels). Sorted cells were seeded on collagen-coated plates and cultured for 24h. Images were taken using a Zeiss fluorescent microscope (right panel). Scale bar: 20μm. **C.** Data analysis workflow. LSEC: liver sinusoidal endothelial cell. **D.** Uniformed Manifold Approximation and Projection (UMAP) showing sorted cell populations in control and cirrhotic mice. The cells were divided into 12 clusters. Each dot represents a single cell. **E.** GFP (left) and Cdh5 (cadherin-5, also known as ve-cadherin; right) expression among the sorted cells. All the cells used for analyses were positive with GFP and Cdh5, indicating a high purity of liver EC populations. **F.** UMAP showing 12 identified clusters of sorted cells. The identity of each cluster was determined by matching expression profiles of clusters with established cell-specific marker genes of different hepatic cells, including ECs, lymphatic ECs, hepatic stellate cells (HSCs), Kupffer cells (KCs), hepatocytes (Hepts) and cholangiocytes. Numbers in parentheses indicate corresponding clusters. **G.** Heatmap showing representative genes expressed by each liver EC cluster/population.

### Spatial lobular locations of heterogeneous liver EC populations were determined

Before comparing transcriptomic differences and related biological changes in each EC cluster/population between control and cirrhotic mice, we first determined a spatial distribution of each cluster in the control (normal) mouse liver based on expression of well-known landmark genes^(1 7–9)^. Consistent with a previous study^(1 10)^, most of the EC genes analyzed exhibited spatial gradations rather than binary expression patterns without clear boundaries between different EC clusters except for Cluster 6.

#### Clusters 1-5 (EC1 – EC5): An atlas of LSECs and vascular ECs (i.e., arterial and central venous ECs) in the control mouse liver

*Because LSECs are unique ECs, we differentiated LSECs from vascular ECs such as arterial* and central venous ECs, using currently known vascular and LSEC makers in Clusters 1 through 5. We found expression of a vascular EC marker, vWF, was much higher in Clusters 1 and 5 than Clusters 2, 3 and 4. In contrast, an LSEC marker, Lyve1 was expressed at a higher level in Clusters 2, 3 and 4 than Clusters 1 and 5 (Figure 2A). In addition, these three clusters expressed other LSEC markers, such as Cd32b, Flt4 and Stab2, at much higher levels than Clusters 1 and 5. Cd31 (a.k.a., Pecam1) has been reported to be more highly expressed in vascular ECs than LSECs and has been used as a marker of capillarization^(11–13)^. However, our data showed that all liver ECs expressed Cd31 with a slightly higher expression in Clusters 1 and 5 than Clusters 2, 3 and 4 (Figure 2A). Collectively, these results indicate that Clusters 1 and 5 likely represent vascular EC populations, while Clusters 2, 3 and 4 correspond to LSEC populations.

**Figure 2.**
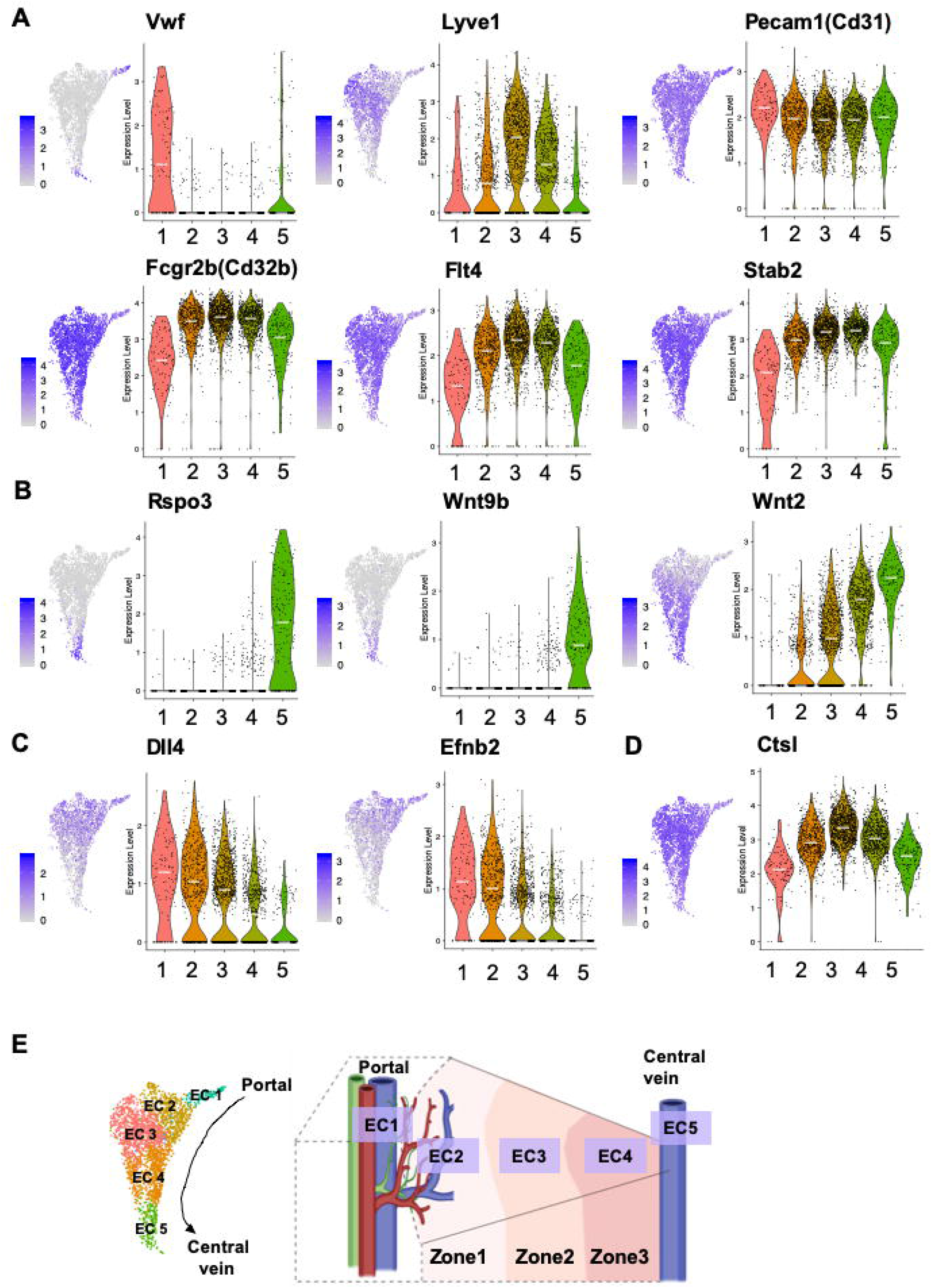
Spatial distributions of the identified liver EC populations (Clusters 1-5) in control mice. **A-D.** Paired feature plots (left) and violin plots (right) showing expression levels of vascular endothelial cell (EC) and liver sinusoid endothelial cell (LSEC) marker genes **(A)**, pericentral landmark genes **(B)**, periportal landmark genes **(C)** and mid-zonal landmark genes **(D)** among liver ECs 1-5. Each dot represents a single cell. In the violin plots, white lines indicate median expression values. **E.** The identified liver ECs 1-5 were mapped on the liver lobule based on expression levels of the marker genes analyzed above and were defined as vascular ECs (EC1), periportal (Zone1) LSECs (EC2), mid-zonal (Zone2) LSECs (EC3), pericentral (Zone3) LSECs (EC4), and central venous ECs (EC5).

It was reported that Rspo3, Wnt9b and Wnt2 were enriched in central venous ECs^(8 9)^. A recent paired-cell sequencing study showed these genes to be pericentral landmarks of liver ECs^(1)^. In our study, Rspo3 and Wnt9b were specifically expressed in Cluster 5, while Wnt2 expression increased gradually from Clusters 2 to 5 with the highest expression in Cluster 5 (Figure 2B). In addition, we found that other pericentral landmarks, such as Kit, Cdh13, Thbd and Fabp4^(1)^, exhibited expression patterns similar to that of Wnt2 (Supplemental Figure 2A). Based on these observations, we consider Clusters 4 and 5 to be a pericentral LSEC population (i.e., Zone3 LSECs) and a central venous EC population, respectively.

We then examined expression patterns of periportal landmarks, such as Dll4 and Efnb2^(1)^. They were also reported to be highly expressed in arterial ECs^(7 14)^. Our analysis showed expression of Dll4 and Efnb2 were both the highest in Cluster 1 with gradual decreases toward Cluster 5 (Figure 2C). Other periportal landmarks, such as Msr1, Ltbp4, Ntn4 and Adam23^(1)^, also showed similar patterns to DII4 and Efnb2 (Supplemental Figure 2B). These results led us to define Cluster 1 as an arterial EC (or portal EC) population and Cluster 2 as a periportal LSEC population (i.e., Zone1 LSECs). Accordingly, Cluster 3 was thought to consist of mid-zonal (Zone 2) LSECs, characterized by the highest expression of mid-zonal landmarks, Lyve1 and Ctsl ^(1 15)^(Figures 2A&D). These results indicated that ECs of Clusters 1 - 5 aligned from the portal tract to the central venous regions as shown in Figure 2E.

We also examined functional differences between periportal (Zone 1) and pericentral (Zone 3) LSECs using Gene Set Enrichment Analysis (GSEA) and identified distinct signaling pathways in Zone1 and Zone3 LSECs (Supplemental Figure 2C). Given that zonal changes are gradual, we expected that LSECs of Zones 1 and 3 might reveal clearer differences in pathways than those of Zones 1 and 2 or Zones 2 and 3. Zone1 LSECs showed a high expression of genes related to netrin-1 signaling, EPH-ephrin mediated repulsion of cells, and antigen processing-cross presentation. Zone3 LSECs were enriched for pathways related to platelet activation signaling/aggregation, hemostasis, and WNT signaling.

#### Cluster 6: Lymphatic ECs

Cluster 6 was identified as lymphatic ECs based on the expression of four well-known lymphatic EC markers, Lyve1, Flt4, Pdpn and Prox1 (Figure 3A). It is known that LSECs also express Lyve1 and Flt4, which were more highly expressed in Zone2 LSECs than LSECs of any other zones in our analysis (Figure 2A). Therefore, we specifically compared expression levels of these four lymphatic EC markers between Zone2 LSECs and lymphatic ECs (Figure 3B). Lyve1 expression was higher in lymphatic ECs than in Zone2 LSECs, while Flt4 expression was similar between these two groups of ECs. Pdpn and Prox1 were specifically expressed in lymphatic ECs. We also identified additional genes that were highly expressed in lymphatic ECs, but not in LSECs, including Mmrn1, Rassf9, Tbx1 and Ahnak2 (Figure 3C), and confirmed their expression by qPCR using primary human LSECs and LyECs (Figure 3D).

**Figure 3.**
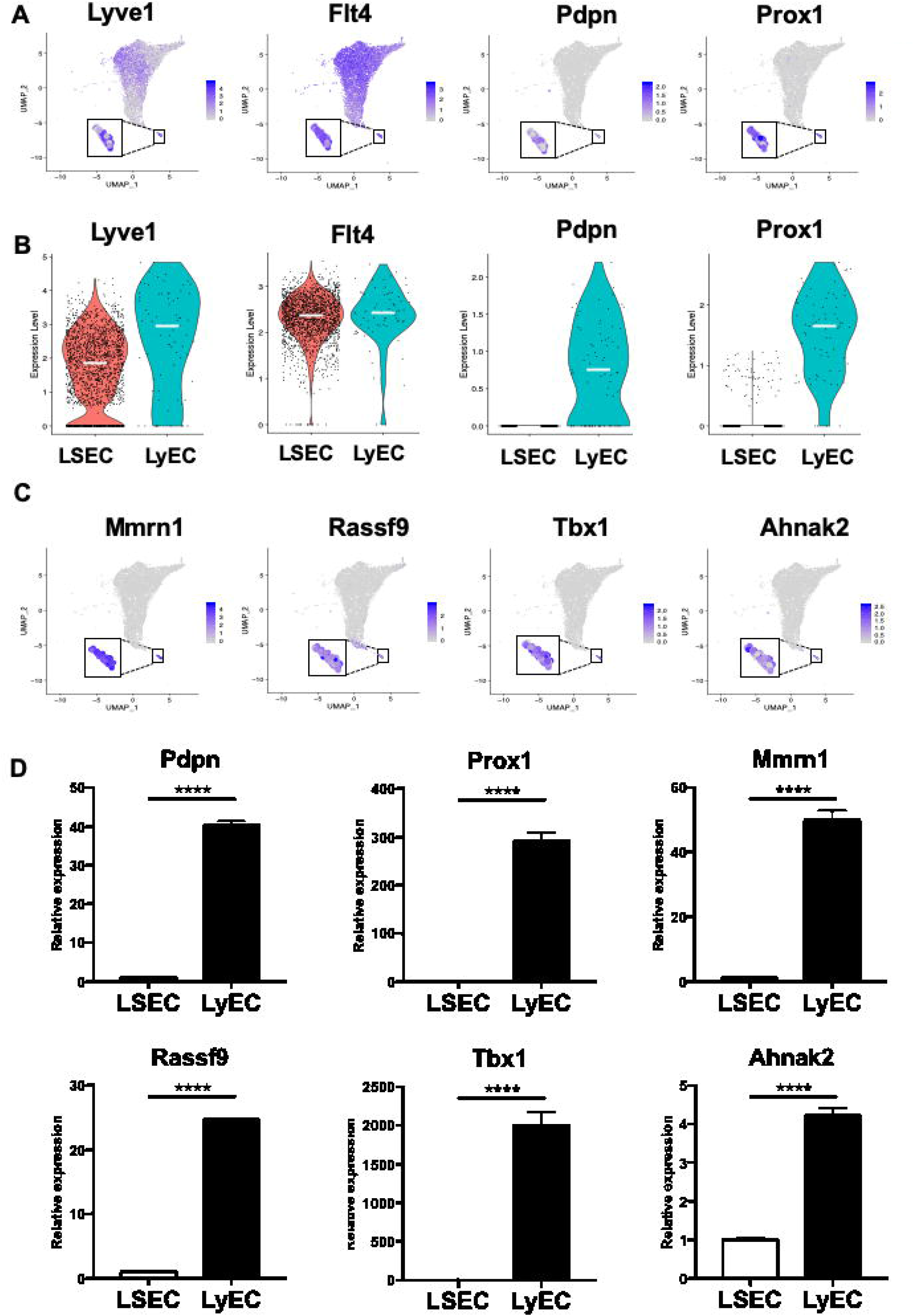
Cluster 6 represents lymphatic EC. **A.** Feature plots showing relative distributions of established lymphatic EC marker genes (Lyve1, Flt4, Pdpn and Prox1) among all the liver ECs. Expression levels of these lymphatic EC marker genes identified Cluster 6 as lymphatic ECs. **B.** Comparison of lymphatic EC maker gene expression between Zone2 LSECs and lymphatic ECs (LyECs). Zone2 LSECs were chosen for the comparison because of the highest levels of Lyve1 and Flt4 they expressed among three LSEC populations. Each dot represents a single cell. White lines indicate median expression values. **C.** Feature plots showing some of the genes found only in Cluster 6, which thus have the potential as new lymphatic EC markers and could help to distinguish LyECs from LSECs. **D.** qPCR analysis to validate unique LyEC markers (distinct from LSECs) identified in this scRNA-seq analysis. Human primary LSECs and LyECs were used for qPCR analysis. n=3. ****p<0.0001. qPCR analysis was repeated three times to confirm this finding.

### EC subtypes in the entire liver EC population in control vs. cirrhotic mice

LSECs accounted for the major portion of the entire liver EC population in both control and cirrhotic mice with 89% and 73%, respectively (Figure 4A). However, the proportions of vascular ECs (Clusters 1 and 5) increased by 2 to 3 times in cirrhotic mice, possibly related to increased angiogenesis in cirrhotic livers^(16)^. Lymphatic ECs represented only 0.12% of all liver ECs in control mice, but increased by 20-fold to 2.34% in cirrhotic mice, which was validated in immunofluorescence images of lymphatic vessels in cirrhotic and control livers (Figure 4B). Interestingly, although cirrhosis changed the proportions of these EC subtypes, spatial EC landmark genes were well conserved between control and cirrhotic livers (Figure 4C).

**Figure 4.**
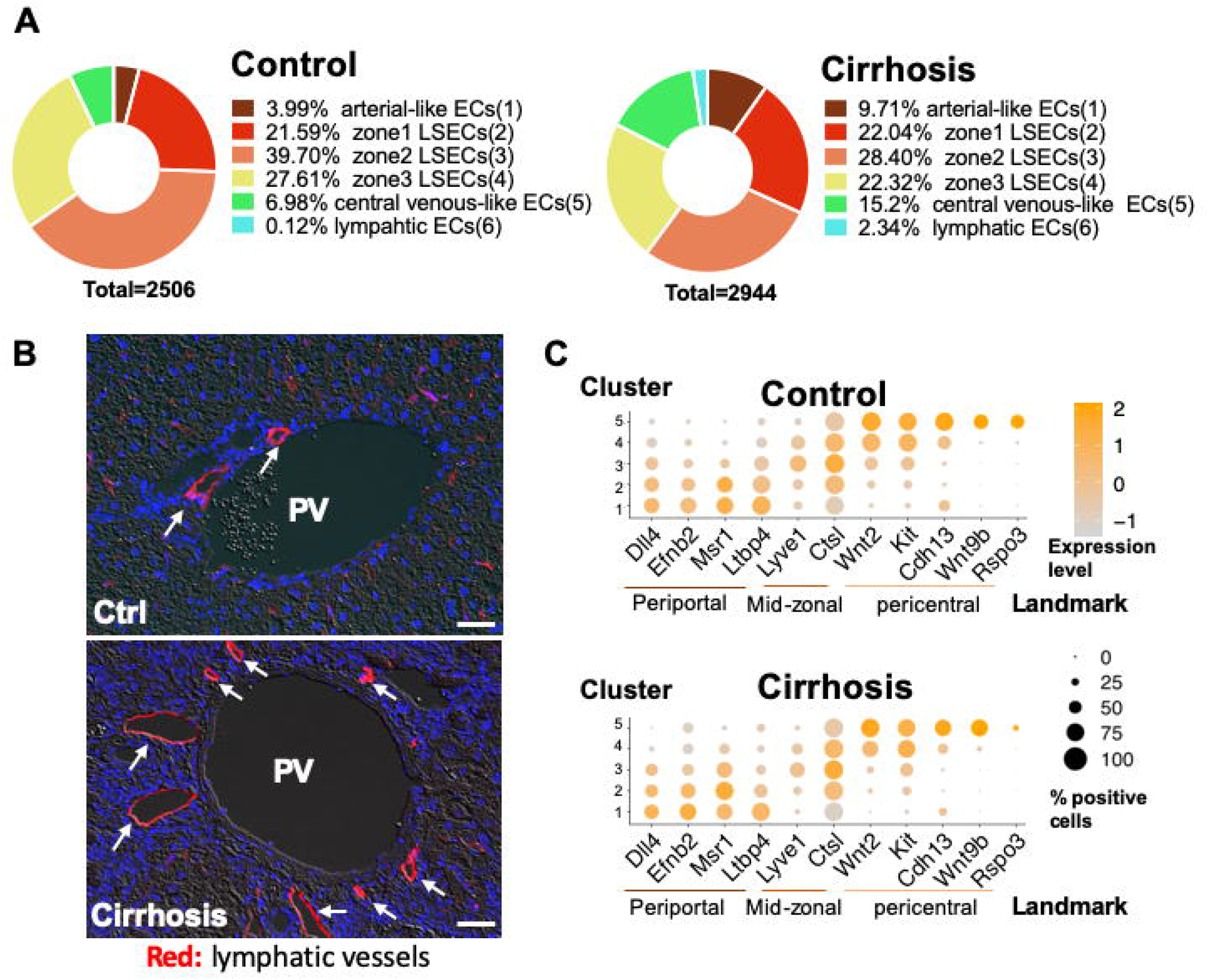
Liver cirrhosis alters proportions of liver EC populations, but still conserves their identities. **A.** Proportions of liver EC populations in control versus cirrhotic mice. **B.** Immunofluorescence images of lymphatic vessels (arrows) in control and cirrhotic mouse livers. Red: Lyve1 (arrows: lymphatic vessels), Blue: DAPI (a marker of nuclei). Because it has been known that a majority of lymphactic vessels are found in the portal tract area, Lyve1 can still be used as a lymphatic vessel marker. PV: portal vein. Scale bar: 40μm. Images were taken using a Zeiss fluorescence microscope. **C.** Dot plots showing conserved landmarks of liver ECs in control and cirrhotic mice. Y-axis indicates EC populations corresponding to Clusters 1- 5 and X-axis refers to EC landmarks, including periportal, mid-zonal and pericentral landmarks. The size of each dot represents a percentage of cells that positively express the landmark gene. Orange color indicates higher expression levels while grey color depicts lower expression levels.

### Phenotypic and functional changes of LSECs in cirrhotic livers

Since LSECs account for the majority of the entire liver EC population, we examined transcriptomic changes in LSECs in liver cirrhosis. In particular, we related spatial changes in gene expression to phenotypic alterations in LSECs. We also re-evaluated representative markers associated with these phenotypic alterations of LSECs.

#### Capillarization was most prominent in Zone3 LSECs and represented by CD34 induction in cirrhotic livers

Capillarization of LSECs is characterized by their phenotypic changes towards common vascular ECs, which is known to cause activation of hepatic stellate cells (HSCs) and thereby liver fibrosis/cirrhosis progression^(17)^. Comparison of gene expression associated with LSEC capillarization between control and cirrhotic livers revealed downregulation of LSEC markers such as Lyve1, Cd32b and Flt4 in cirrhotic mice (Figure 5A). Many studies have used upregulation of CD34 and/or CD31 in LSECs as a sign of LSEC capillarization^(11 12 18 19)^. We found significant upregulation of Cd34 in all zones of LSECs of cirrhotic mice (average fold change=6.3) with its very low expression in LSECs of control mice (Figures 5B&F), which was consistent with immunolabeling results in Figure 5C (left panels), showing prominent expression of CD34 around Zone 3 in cirrhotic liver, but very low expression in control liver. In contrast, CD31 was highly expressed in LSECs regardless of the presence of cirrhosis with only a slight upregulation in cirrhotic liver (average fold change=1.1) (Figure 5B, Supplemental Figure 3B), which was also verified by immunolabeling results (Figure 5C right panels, Supplemental Figure 3A). These results indicate that CD34 is a more accurate marker of LSEC capillarization than CD31.

**Figure 5.**
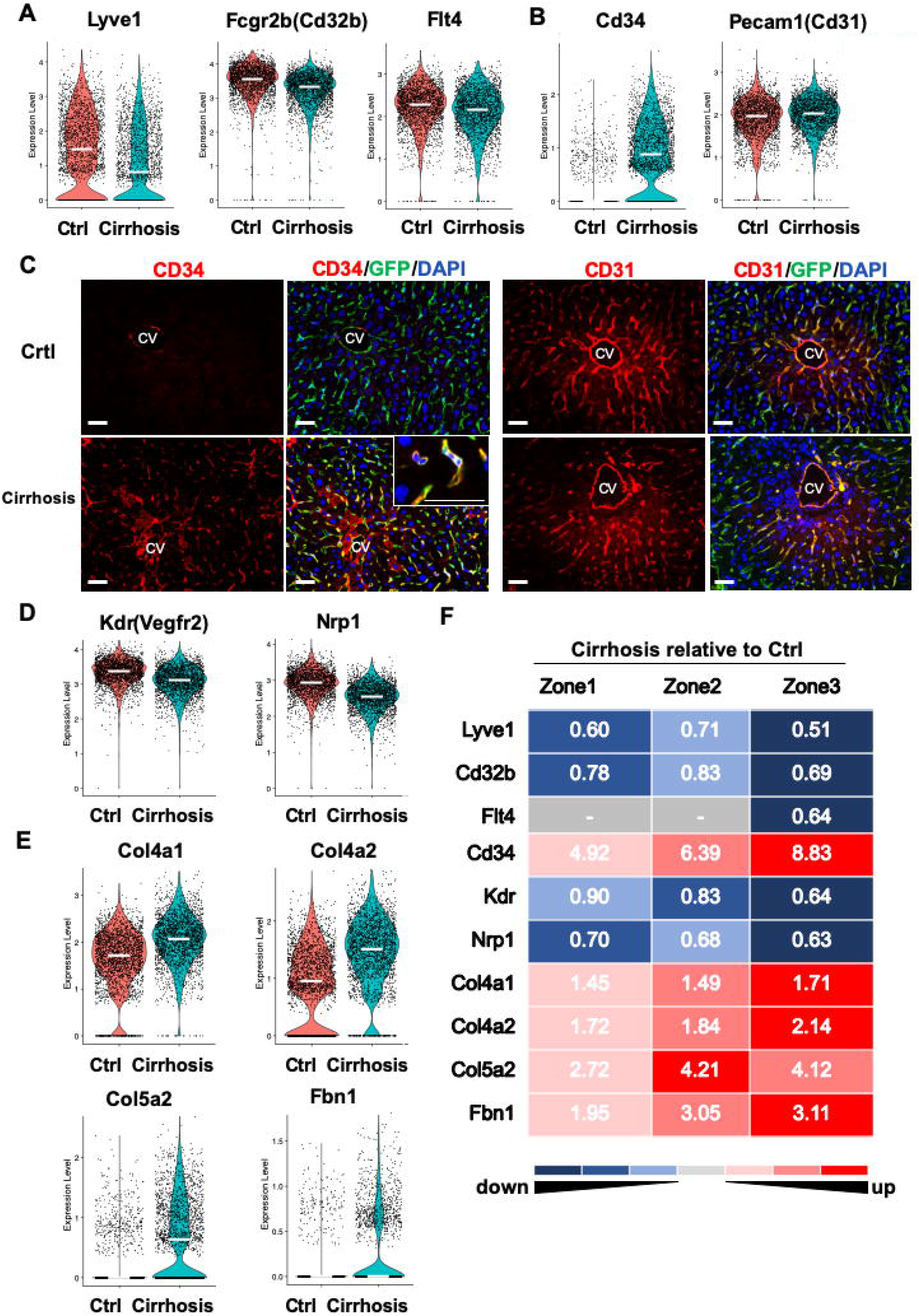
LSEC capillarization is most prominent in Zone 3 and CD34 represents LSEC capillarization more accurately than CD31. **A&B.** Expression of capillarization-associated genes in LSECs (Clusters 2, 3 and 4) of control and cirrhotic mice. Each dot represents a single cell. White lines indicate median expression values. **C.** Immunofluorescence staining of CD34 or CD31 in frozen liver tissue sections from endothelial-GFP reporter mice. Red: CD34 or CD31 (frequently used capillarization markers), Green: GFP (endothelial cells), Blue: DAPI (nuclei). CV: central vein. Scale bar: 40μm. Images were taken using a Zeiss fluorescence microscope. **D&E.** Expression of Kdr, Nrp1 (genes to maintain LSEC phenotype) **(D)** and extracellular matrix (ECM) genes **(E)** in LSECs (Clusters 2, 3 and 4) of control and cirrhotic mice. Each dot represents a single cell. White lines indicate median expression values. **F.** Differential gene expression between control and cirrhotic mice in each cluster of LSECs (corresponding to Zone 1, 2 or 3). Red indicates upregulation, while blue means downregulation in cirrhotic mice compared to control mice with the numbers indicating fold changes (cirrhosis relative to control). The darker the color, the more profound the changes of gene expression. Gray indicates no significant differences.

Previous studies also reported that VEGF released by hepatocytes and HSCs maintained LSEC phenotype in a paracrine manner^(20)^. We found a VEGF receptor, Kdr (a.k.a., Vegfr2), and its co-receptor Nrp1 were both downregulated in LSECs of cirrhotic mice (Figure 5D), which may also explain decreased VEGF signaling and subsequent dysregulation of LSEC phenotype in cirrhotic livers. In addition, we found that extracellular matrix (ECM) genes, such as Col4a1, Col4a2, Col5a2 and Fbn1, were all upregulated in LSECs of cirrhotic mice (Figure 5E), which may be related to the development of basement membranes (typical of LSEC capillarization) and ECM deposition in liver fibrosis/cirrhosis. Comparison of zonal expression of the above-mentioned capillarization-associated genes between control and cirrhotic mice revealed that almost all these genes were most downregulated or upregulated in Zone3 LSECs (Figure 5F), suggesting Zone3 LSECs are the most susceptible to capillarization in liver cirrhosis.

#### Decreased endocytic capacity

LSECs are involved in removal of circulating antigens and toxins through their strong endocytic capacity^(21)^. We found that expression of major endocytic receptors including mannose receptor (Mrc1), scavenger receptors (Stab1, Stab2, Scarb1 and Scarb2) and lysosome-associated membrane glycoprotein 2 (Lamp2)^(22)^ were significantly decreased in LSECs of cirrhotic mice (Figure 6A). Interestingly, all these endocytosis-associated genes were also most downregulated in Zone3 LSECs of cirrhotic mice (Figure 6B).

**Figure 6.**
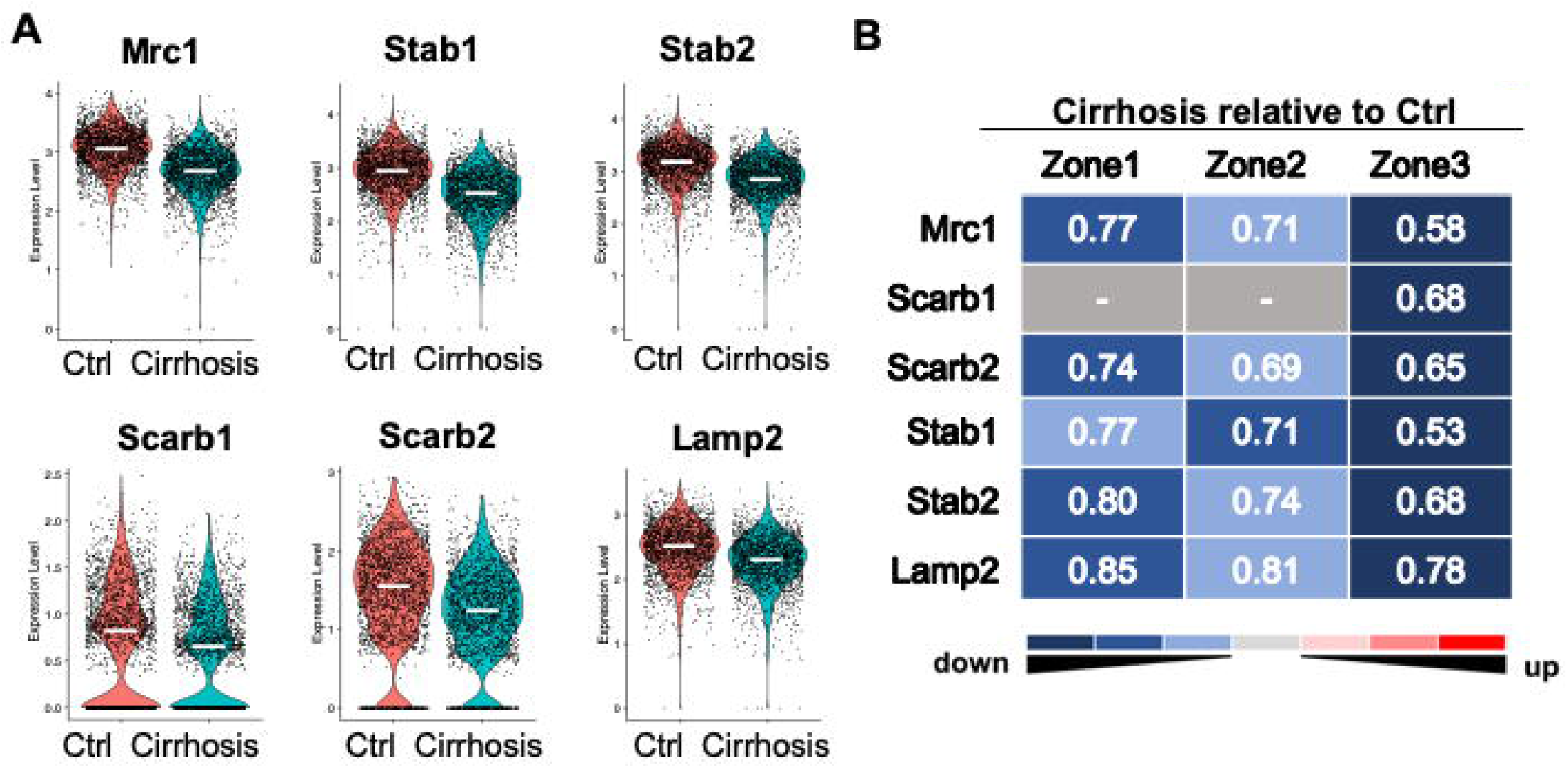
Cirrhosis decreases expression of endocytotic receptor genes most profoundly in Zone3 LSECs. **A.** Expression of endocytosis receptor genes in LSECs (Clusters 2, 3 and 4) of control and cirrhotic mice. Each dot represents a single cell. White lines indicate median expression values. **B.** Differential expression of endocytosis receptor genes between control and cirrhotic mice in each cluster of LSECs (corresponding to Zone 1, 2 or 3). Blue means downregulation in cirrhotic mice compared to control mice with the numbers indicating fold changes (cirrhosis relative to control). The darker the color, the more profound the changes of gene expression. Gray indicates no significant differences.

#### Regulation of vascular tone

LSECs respond to increased shear stress to maintain normal vascular tone by promoting nitric oxide (NO) production by endothelial NO synthase (eNOS)^(23)^. The loss of this property is one of the representative features of endothelial dysfunction and is observed in cirrhosis ^(21 24)^. Some transcription factors, such as the Kruppel-like family (Klf2 and Klf4) and activating protein-1 (AP1), are induced by shear stress and are responsible for increased eNOS expression and activity^(25–28)^. We found downregulation of both Klf2 and Klf4 in LSECs of cirrhotic mice (Figure 7A, Supplemental Figure 4A). Similarly, some of the major AP1 components, such as Fos, Fosb, Jun and Junb, were remarkably suppressed in LSECs of cirrhotic livers (Figure 7B, Supplemental Figure 4A).

**Figure 7.**
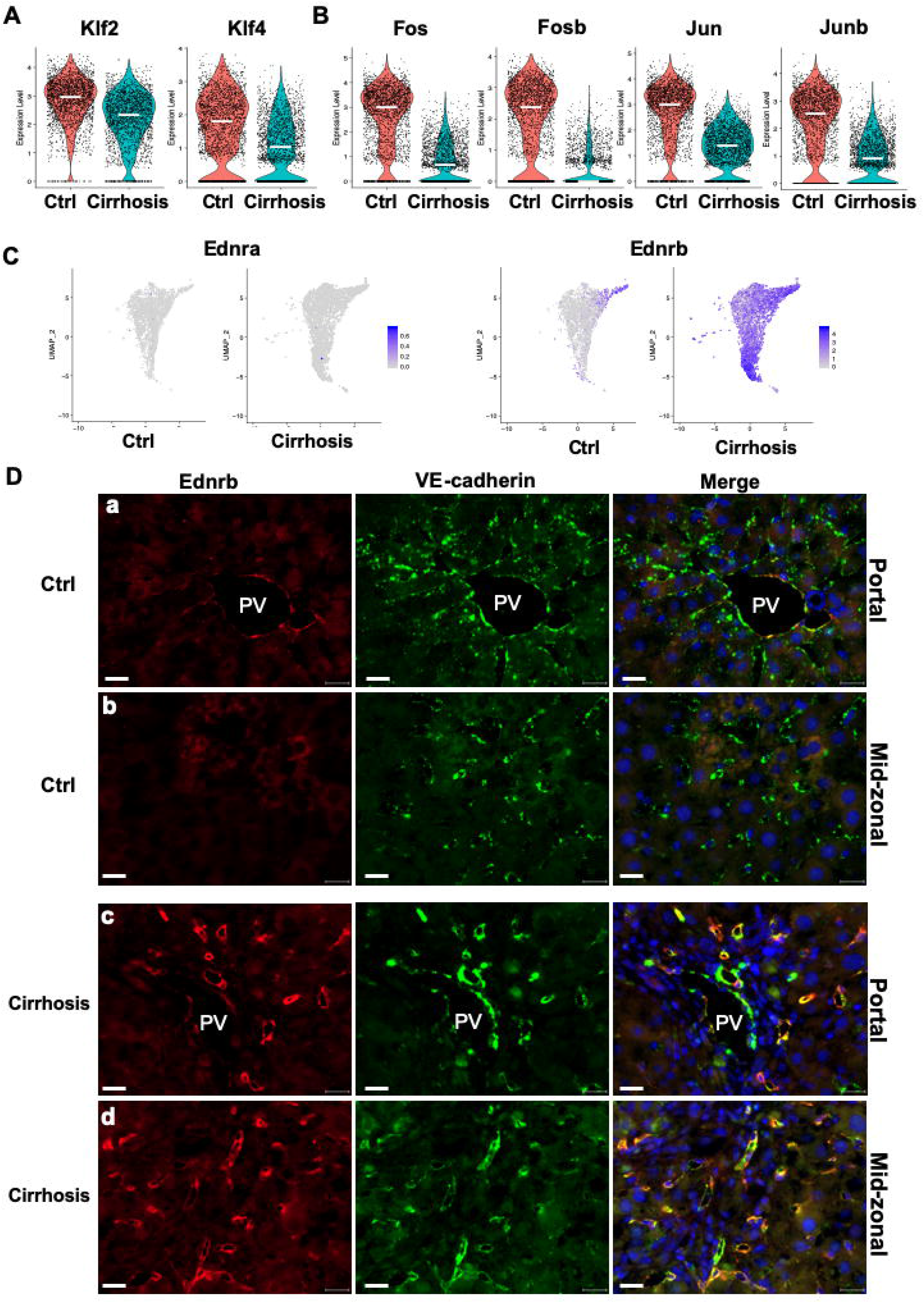
Identification and validation of genes associated with LSEC dysfunction in cirrhotic livers. **A&B.** Expression of transcription factors in LSECs (Clusters 2, 3 and 4) of control and cirrhotic mice. Each dot represents a single cell. White lines indicate median expression values. **C.** Relative distributions of endothelin receptors (Ednra and Ednrb) in liver ECs of control and cirrhotic mice. **D.** Immunofluorescence staining of Ednrb in frozen liver tissue sections from endothelial-GFP reporter mice. The portal tract area (a&c) and the mid-zonal area (b&d) of the liver. Red: Ednrb, Green: VE-cadherin (represents all liver ECs), Blue: DAPI (nuclei). PV: portal vein. Scale bar: 20μm. Images were taken using a Zeiss fluorescence microscope.

One of the key signaling pathways regulating sinusoidal vascular tone is the endothelin signaling pathway. Endothelin receptor type A (ET_A_) and B (ET_B_) are encoded by Ednra and Ednrb genes, respectively^(29)^. In our analysis, Ednra was not expressed in LSECs either in control or cirrhotic livers (Figure 7C left two panels) as it is known to be expressed in smooth muscle cells, not in ECs^(29)^, while Ednrb was significantly upregulated in LSECs of all zones in cirrhotic livers (Figure 7C right two panels, Supplemental Figure 4B). In control livers, Ednrb was highly expressed only in portal ECs and some adjacent LSECs. Immunolabeling of Ednrb in control and cirrhotic livers was consistent with scRNA-seq results (Figure 7D).

#### Endothelial-to-mesenchymal transition (EndMT)

Several studies reported that LSECs undergo EndMT in response to chronic liver injury^(30–32)^. In contrast, we did not find notable increases in mesenchymal markers, such as α-SMA, Sm22, Fn1 and Fsp1, in LSECs of cirrhotic mice compared to those of control mice (Figure 8A). In addition, with the exception of vimentin, other EndMT-associated genes, such as Snail1&2, Twist1, Zeb1&2, Col1a1&2, Tgfb2&3, Tgfbr3 and Tgfbi, were not upregulated in LSECs of cirrhotic mice either (Supplemental Figure 5). The absence of EndMT in LSECs of cirrhotic mice was also demonstrated by immunolabeling of α-SMA in livers from EC-GFP reporter mice subjected to CCl_4_ inhalation for 12 weeks to induce liver cirrhosis (Figure 8B) or bile duct ligation (BDL) for 1, 2 and 4 weeks to induce liver injury (1-week BDL), fibrosis (2-week BDL) and cirrhosis (4-week BDL) (Figure 8C). GFP-positive cells representing all liver ECs did not co-localize with α-SMA in LSECs in either CCl_4_ or BDL models (Figures 8B&C). However, it is noted that in an *in vitro* cell culture condition, rat primary LSECs underwent EndMT in a time dependent manner (Supplemental Figure 6). Collectively, these results suggest that mouse LSECs seem resistant to EndMT in liver cirrhosis *in vivo*.

**Figure 8.**
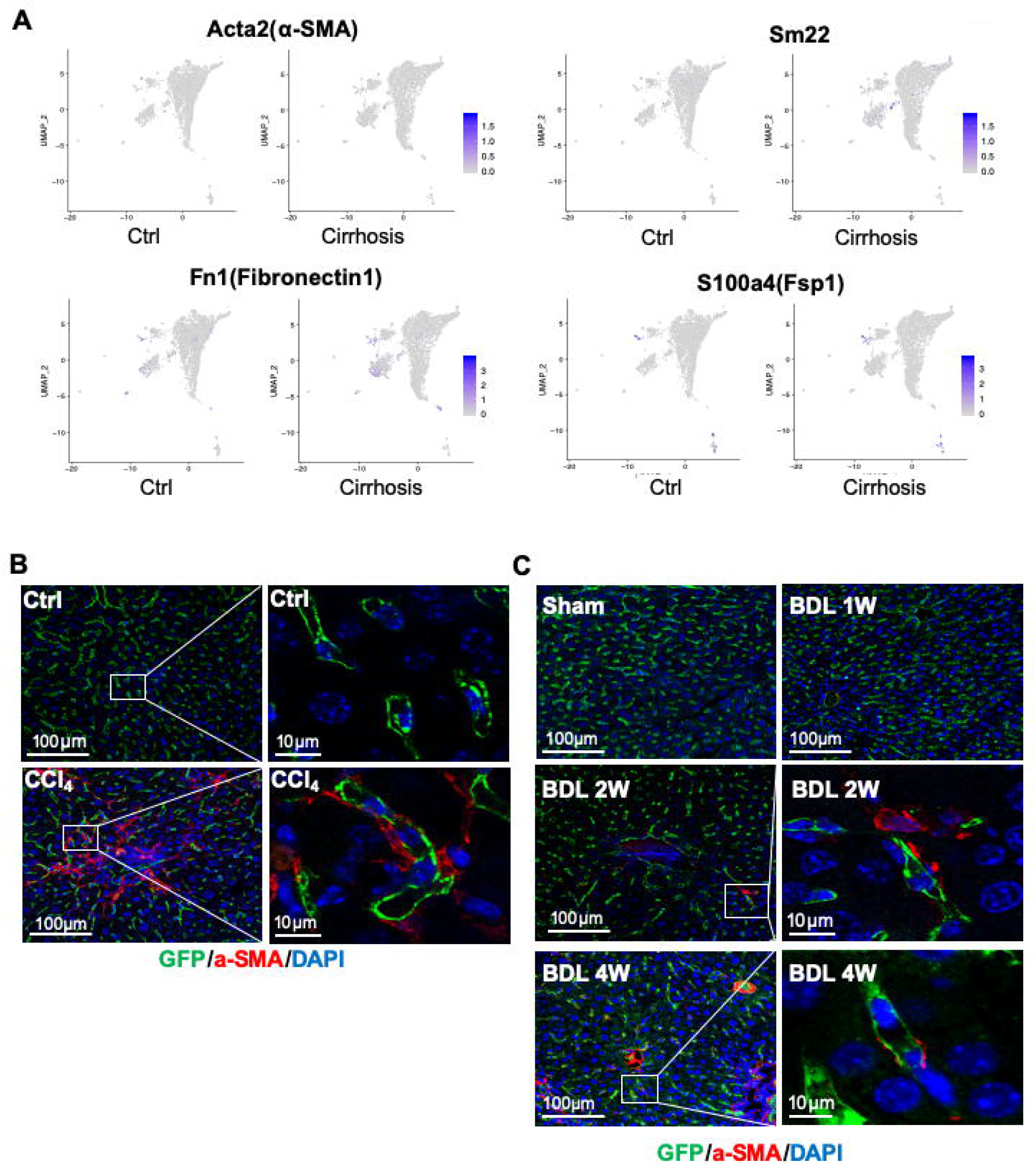
LSECs likely do not undergo endotheial-to-mesenchymal transition (EndMT) in injured, fibrotic and cirrhotic mouse livers. **A.** Relative distributions of mesenchymal marker genes in sorted cells of control and cirrotic mice to evaluate EndMT in liver cirrhosis. **B&C.** Immunofluorescence images of paraffin liver tissue sections (control and fibrotic/cirrhotic livers) isolated from endothelial-GFP reporter mice subjected to CCl_4_ inhalation for 12 weeks (**B**) and bile duct ligation (BDL) (**C**). Green: GFP (endothelial cells), Red: α-smooth muscle actin (α-SMA, a mesenchymal cell marker), Blue: DAPI (nuclei). Images were taken using a confocal fluorescence microscope.

#### Pathway analysis

To examine what biological signaling pathways are potentially altered in whole LSECs or specific zonal LSECs of cirrhotic mice, we performed Gene Set Enrichment Analysis (GSEA). Pathways activated or suppressed in LSECs of cirrhotic mice compared to control mice are presented (Supplemental Figure 7). LSECs in Zones 1, 2 and 3 showed some similarities, but also distinct differences in signaling pathways affected by liver cirrhosis, suggesting zonal specificities of LSEC function. LSECs of all three zones of cirrhotic mice showed upregulation of ribosome, PPAR signaling pathway and downregulation of IL17 signaling pathway. There are some unique pathways only present or absent in specific zones. Such examples found in Zone3 LSECs include: (1) absence of upregulation of rap1 signaling, platelet activation and actin cytoskeleton regulation pathways and (2) presence of upregulation of gap junction signaling pathway.

## Discussion

We identified 6 clusters of mouse liver EC populations, including 3 clusters of LSECs, 2 clusters of vascular ECs and 1 cluster of lymphatic ECs in our scRNA-seq analysis. We also found potential novel lymphatic EC markers distinct from those for LSECs. The particular importance of our study consists in spatial (i.e., zonal) characterization of LSECs (Zones 1–3), identification of transcriptomic changes in these zones associated with liver cirrhosis, and demonstration of relationships between these transcriptomic changes and phenotypic changes observed in liver cirrhosis. We found that heterogeneous liver EC identities are conserved even in liver cirrhosis and that Zone3 LSECs are most susceptible to damages associated with liver cirrhosis with increased capillarization and decreased abilities to regulate endocytosis. Furthermore, we demonstrated that CD34 is more useful as a marker of LSEC capillarization in liver cirrhosis than CD31.

The role of LSECs in the pathogenesis of liver fibrosis/cirrhosis has received a great deal of attention for many years^(21 33)^. Most studies have identified differentially expressed genes by qRT-PCR or bulk RNA-seq using isolated primary LSECs. However, isolation of pure LSECs is challenging, because LSEC preparation can easily be contaminated with other cell types, especially with vascular ECs, which may influence overall interpretation of results. Furthermore, due to heterogeneity of the LSEC population, some isolation techniques may exclude certain subpopulations of LSECs^(5 33)^. For example, Lyve1 is negative in some periportal LSECs resulting in their removal when sorting is based on Lyve1 positivity^(5 34)^. One of the strengths in our study is use of EC-specific GFP reporter mice. Isolating GFP-positive cells from these mice reduced selection bias. In addition, as the isolation did not require extra marker staining, it simplified the isolation process and saved sample preparation time, helping to improve cell viability, which was especially important for fragile cells like LSECs. Furthermore, scRNA-seq analysis based on these isolated GFP-positive cells allowed us to identify highly enriched LSEC populations from all liver ECs and have all subtypes of LSECs. We think that these advantages of cell sorting conferred more reliable and comprehensive qualities to our comparison of differentially expressed genes in these identified LSECs between control and cirrhotic mice.

Capillarization of LSECs, which is characterized by loss of fenestrae and development of basement membranes, is a well-recognized phenotypic change in liver fibrosis/cirrhosis^(21)^. Besides use of electron microscopy for evaluation of fenestrae in LSECs, previous studies have used several markers to indicate LSEC capillarization, such as downregulation of Lyve1 and CD32b and upregulation of CD31 and CD34^(11 13 18 35 36)^. While CD31 is a frequently used maker for LSEC capillarization^(11–13)^, expression of CD31 by LSECs has been controversial. Some studies indicated absence of CD31 in normal LSECs but its presence in LSECs in cirrhosis^(37–39)^. Other studies showed that CD31 is already highly expressed in LSECs in normal livers^(10 40 41)^ and its levels are not changed in liver disease ^(40 41)^. Our scRNA-seq analysis together with immunolabeling analysis provide strong evidence to support the latter observation with similar expression levels of CD31 in LSECs between normal and cirrhotic livers (Figures 5B&C, Supplemental Figure 3). A study using immunogold-scanning electron microscopy showed that CD31 was located intracellularly one day after primary LSECs were cultured and moved to the cell surface a few days later when fenestrae disappeared^(42)^. Changes in the cellular location of CD31 may influence its detection by flow cytometry or immunolabeling. It should also be noted that the proportion of vascular ECs increases in liver cirrhosis as angiogenesis increases^(43)^. This was also confirmed by our scRNA-seq analysis (Figure 4A). Contamination of vascular ECs in isolated primary LSECs could thus lead to overestimation of CD31 expression by LSECs in cirrhotic livers. Therefore, CD31 may not be an appropriate marker for LSEC capillarization if its expression levels are solely assessed without any consideration of its cellular location. In contrast, CD34 is barely expressed in normal LSECs, but highly expressed in LSECs in cirrhotic livers, suggesting that CD34 is a reliable indicator of LSEC capillarization.

We found that capillarization was most severe in Zone3 LSECs, suggesting pericentral LSECs are most vulnerable in the microenvironment of cirrhotic livers. Since blood runs from portal veins and hepatic arteries towards central veins, creating decreasing gradients of oxygen and nutrition along liver lobules with their lowest levels in the central vein areas, hepatocytes in the pericentral area may be more sensitive to anoxia and damage in cirrhotic livers. An interaction of injured hepatocytes and LSECs in Zone 3 may aggravate capillarization of LSECs. The mechanism of LSEC capillarization is still not well understood. It is reported that VEGF produced by hepatocytes and hepatic stellate cells maintain the phenotype of LSECs^(20)^. However, VEGF secretion is increased in cirrhotic livers^(21)^, suggesting that capillarization of LSECs may be related to disruption of downstream signaling of VEGF rather than lack of VEGF. We found both VEGF receptor Kdr and co-receptor Nrp1 were most downregulated in Zone3 LSECs of cirrhotic mice as well, which may contribute to LSEC capillarization to some degree.

LSECs are one of the most powerful scavengers in the body, playing an important role in clearance of wastes and pathogens in blood originated from the gut and the systemic circulation ^(44–47)^. This activity is related to their expression of various endocytosis receptors including scavenger receptors (Scarb1, Scarb2, Stab1 and Stab2) ^(47)^, mannose receptor (Mrc1) ^(48)^ and Fc gamma-receptor IIb2 (Fcgr2b/CD32b)^(49)^. We found downregulation of all these endocytosis receptors in cirrhotic livers, suggesting decreased endocytic and clearance capacities of LSECs (Figure 6). This may make cirrhotic patients more susceptible to infection and systemic inflammation. Interestingly, all these endocytosis receptors were also most downregulated in Zone3 LSECs in cirrhotic mice. The decreased endocytic capacity of LSECs may be associated with their capillarization as well, because decreased CD32b was also used as an indicator of LSEC capillarization in some studies ^(19 50 51)^. Identification of the most dysfunctional LSEC populations will be tremendously useful for the development of effective therapeutic strategies targeting them.

We also found that genes known to promote eNOS expression were downregulated in LSECs of cirrhotic mice, indicating dysfunction of vascular tone observed in cirrhosis. However, our analysis also showed Ednrb expression (ET_B_ receptor), known to increase NO signaling in ECs^(29)^, was upregulated in cirrhotic livers. Ednrb upregulation was reported in human cirrhotic livers at both mRNA and protein levels as well^(52 53)^. Upregulation of Ednrb in LSECs could be an adaptive response to compensate for the loss of NO signaling in cirrhotic livers. Or, endothelial ET_B_ receptor may have different activities in physiological versus pathological conditions. One study reported that endothelial ET_B_ receptor contributed to vasodilatation in healthy vessels, but that endothelial ET_B-_mediated vasodilatation was lost in rats with pulmonary or systemic hypertension and turned into vasoconstriction^(54)^. In patients with cardiovascular pathologies such as atherosclerosis and/or type 2 diabetes, ET_B-_mediated vasodilatation is also lost^(55 56)^. It was reported endothelin-1 could increase expression and activity of arginase-2 ^(57)^ as well as oxLDL receptor-1 (LOX1) ^(58)^ via endothelial ET_B_ receptor in atherosclerotic disease. Arginase-2 can reduce NO production by competing with eNOS for a common substrate (L-arginine)^(57)^, while oxLDL is able to impair endothelial relaxation by reducing eNOS expression and inducing reactive oxygen species^(58)^. Further, chronic ET_B_ antagonism in cirrhosis was shown to lead to less fibrosis^(59)^. This may suggest that overexpression of ET_B_ receptor by LSECs may have a profibrotic effect. Thus, it is possible that ET_B_ receptor in LSECs of cirrhotic livers may have other dominant downstream signaling pathways associated with endothelial dysfunction, which is an important area of future research.

Endothelial-to-mesenchymal transition (EndMT) refers to a process whereby ECs lose endothelial markers like VE-cadherin and CD31 and gain mesenchymal markers such as α-SMA and Fsp1^(60)^. Several studies on fibrotic diseases, including cardiac fibrosis^(61)^, renal fibrosis ^(62)^and pulmonary fibrosis^(60 63)^, indicated that ECs could give rise to myofibroblasts through EndMT. In liver fibrosis too, some studies indicated EndMT in LSECs ^(30–32)^. In contrast, we did not find evidence of EndMT in LSECs in cirrhotic livers in our scRNA-seq data as well as immunostaining of liver sections from two models of liver fibrosis/cirrhosis [CCl_4_ inhalation and bile duct ligation (BDL) models] using EC-GFP reporter mice (Figure 8), although LSECs underwent EndMT in a cultured condition in a time dependent manner (Supplemental Figure 6). Even if liver ECs underwent EndMT *in vivo*, their population would be very small, as also indicated by Ribera et al.^(30)^ who observed EndMT only in about 4% of the liver EC population from cirrhotic livers. Our results indicate that unlike ECs in other organs, LSECs seem highly resistant to EndMT even in conditions of chronic stress and injury. Identification of the mechanism preventing LSECs from EndMT *in vivo*, but not *in vitro*, may help to develop and/or maintain LSECs that can be used for a variety of research and clinical purposes including generation of an engineered liver.

A recent scRNA-seq study of human-derived liver nonparenchymal cells (NPCs) from normal and cirrhotic patients identified two disease-specific EC populations, characterized by CD34^+^PLVAP^+^VWA1^+^ and CD34^+^ PLVAP^+^ACKR1^+(6)^. The authors named them “scar-associated ECs”, but did not demonstrate their origins. Our study did not find any disease-specific EC populations and showed similar genetic landscapes of liver ECs between control and cirrhotic mice as demonstrated in (Figures 1D&4C). However, similar to the study of NPCs from cirrhotic patients, we observed significant upregulation of CD34 (Supplemental Figure 3), PLVAP and ACKR1 (Supplemental file of differentially expressed genes) in LSECs of all zones in cirrhotic livers compared to control livers. This result may suggest that the disease-specific EC populations found in human cirrhotic livers derive from LSECs, whose gene expression profiles are altered in liver cirrhosis. The presence of the disease-specific EC populations might also be attributable to the heterogeneity of genetic backgrounds and/or different stages of liver fibrosis in those human patients. Otherwise, the difference between their results and ours may come from the difference of the study subjects, i.e., humans and mice. It should be mentioned that the same group of researchers recently showed the zonation pattern of hepatic stellate cells (HSCs), which was conserved between healthy and fibrotic mouse livers ^(3)^. Interestingly, they also found that peri-central HSCs were predominant pathogenic collagen-producing HSCs in liver fibrosis, which may be related to our finding that Zone3 LSECs are most susceptible to capillarization.

Of note, we employed a 12-week CCl_4_ inhalation model for our scRNA-seq analysis, which generated end stage liver cirrhosis in mice. It would be of interest to monitor transcriptomic changes in LSECs during the course of liver fibrogenesis, which can further advance our understanding of temporal transcriptomic changes that lead to LSEC dysfunction and their relationship with the development of fibrosis/cirrhosis.

In conclusion, the current study illustrated zonal transcriptomic alterations of LSECs in cirrhotic mouse livers and related them to phenotypic changes of LSECs observed in liver cirrhosis, which deepens our knowledge of the pathogenesis and pathophysiology of cirrhosis at a spatial, cell-specific level and helps to advance biomedical research, both basic and clinical, on liver cirrhosis. In the era of precision medicine, microenvironmental information like that presented in this study is indispensable for the development of novel and effective therapeutic strategies to target the most dysfunctional ECs and mitigate their pro-fibrotic activities in liver fibrosis/cirrhosis^(12)^.

## Materials and Methods

### Animals

Cdh5-CreERT2, mT/mG mice were used ^(64)^. GFP expression in endothelial cells (ECs) was induced by intraperitoneal injection of tamoxifen (Sigma-Aldrich, St. Louis, MO) at a dose of 100μg/g body weight for 5 consecutive days. Liver fibrosis/cirrhosis was induced by inhalation of vaporized carbon tetrachloride (CCl_4_) for 12 weeks ^(65)^ and bile duct ligation (BDL) for 1, 2 or 4 weeks^(66)^. For the CCl_4_ model, mice started to receive the treatment around 4 weeks of age. For the BDL model, mice at the age of around 2 months were used. Age-matched mice and sham-operated mice were used as controls for the CCl_4_ and BDL models, respectively. Animals were allowed free access to food and water and maintained in a 12:12-h light-dark cycle in a temperature (18-21°C) and humidity (55±5%) controlled environment. All animal experiments were approved by the Institutional Animal Care and Use Committees of Yale University and the Veterans Affairs Connecticut Healthcare System and were performed in accordance with the National Institutes of Health Guide for the Care and Use of Laboratory Animals.

### Cell isolation

Liver non-parenchymal cells (NPCs) were isolated from control mice and mice subjected to CCl_4_ inhalation for 12 weeks as previously described with some modifications^(67)^. Briefly, liver cell suspensions were obtained by collagenase perfusion and were spun down at 100 xg for 5 min to remove hepatocytes. The supernatants were pelleted at 350 xg for 10 min and resuspended in Endothelial Cell Growth Basal Medium-2 (EBM-2, CC-3156, Lonza, Morristown, NJ). Isolated NPCs were stained with SYTOX® Red (5uM, Invitrogen, Carlsbad, CA) to label dead cells and sent to FACS sorting. Only live GFP-positive cells were sorted with BD FACSAria™ IIu (BD Biosciences, San Jose, CA) using a 100-μm nozzle.

### 10x sample processing and cDNA library preparation

Samples were prepared according to the instructions of 10x Genomics Single Cell 3, Reagent Kits v3 (10x Genomics, Pleasanton, CA). Briefly, sorted cells (live GFP-positive cells) were pelleted and resuspend to attain a concentration of 1000 cells/ul. The cell number and viability were evaluated again with Trypan Blue (Gibco) and a hemocytometer and confirmed with an automated cell counter. Both samples (control and CCl_4_ groups) consisted of >80% viable cells. Single cell suspensions in RT Master Mix (10x Genomics) were then loaded onto the 10x Genomics Single Cell A Chip to convert poly-adenylated mRNA into barcoded cDNA. Barcoded cDNA was amplified by PCR to generate a sufficient mass for library construction. Enzymatic Fragmentation and Size Selection were then used to optimize the cDNA amplicon size prior to library construction, which included end-repair, A-tailing, adaptor-Ligation, and sample indexing PCR to produces Illumina-ready sequencing libraries.

### Sequencing and data analysis

Sequencing was run on the HiSeq 4000 system (Illumina, San Diego, CA) at the Yale Center for Genome Analysis (YCGA) at Yale University. Each sample was sequenced across 2 lanes of the HiSeq, generating 100bp paired-end reads at a depth of 9000 reads per cell. Preliminary standard analysis steps such as alignment and gene counting were performed based on Cell Ranger™ pipelines (10× Genomics). Cell Ranger outputs were loaded into Seurat v3.0 package (http://satijalab.org/seurat/) to cluster and visualize scRNA-seq data. Genes detected in at least 3 cells were included. Cells that expressed fewer than 200 genes or had high mitochondrial genome transcript ratios (>0.2) were excluded. In order to exclude non-endothelial cells, we filtered out cells that did not express GFP. After normalizing the data using a global-scaling normalization method “LogNormalized” and scaling the data, principal component analysis (PCA) was performed to reduce the number of dimensions. Cell clusters were visualized by Uniform Manifold Approximation and Projection (UMAP). Symbol gene IDs were converted to Entrez gene IDs and gene set enrichment analysis was performed using the ClusterProfilter package^(68)^.

### LSEC and LyEC culture

Refer to the Supplemental Materials.

### Quantitative real-time polymerase chain reaction

Refer to the Supplemental Materials.

### Immunofluorescence

Refer to the Supplemental Materials.

### Statistical analysis

For single-cell RNA seq data, differential expression of genes between clusters or treatment groups were calculated using the Wilcoxon Rank Sum test implemented in Seurat v3.0 package. qPCR results for validation of differentially exressed genes between primary LSECs and lymphatic ECs were evaluated by Student’s t-test using GraphPad Prism 7. Adjusted *p* values or *p* values less than 0.05 were considered statistically significant.

### Data availability

The authors declare that all other data supporting the findings of this study are available within the article and its supplementary information files or from the corresponding author upon request.

## Acknowledgements

This work was funded by NIH grants (R01DK117597 and R56DK121511) to YI; postgraduate fellowship from the Chinese Scholarship Council to TS, YY and SL; AASLD Clinical, Translational, and Outcomes Research Award and NIH grant (T32 DK007356) to MM. We would like to thank Drs. William Sessa and Xinbo Zhang for valuable discussion for the manuscript.

## Author contributions

YI designed and supervised the study, and wrote the manuscript. TS performed experiments, analyzed data and wrote the manuscript. YY, SL, JJ, and YJ performed experiments and analyzed data. MM critically reviewed and wrote the manuscript. TU supervised animal studies, performed all animal experiments, contributed to the design of the study, and critically revised the manuscript. All authors read and approved the final manuscript.

