## Supplemental Materials for "Single-cell transcriptomics reveals zone-specific alterations of liver sinusoidal endothelial cells in cirrhosis"

### Supplemental Fig 1

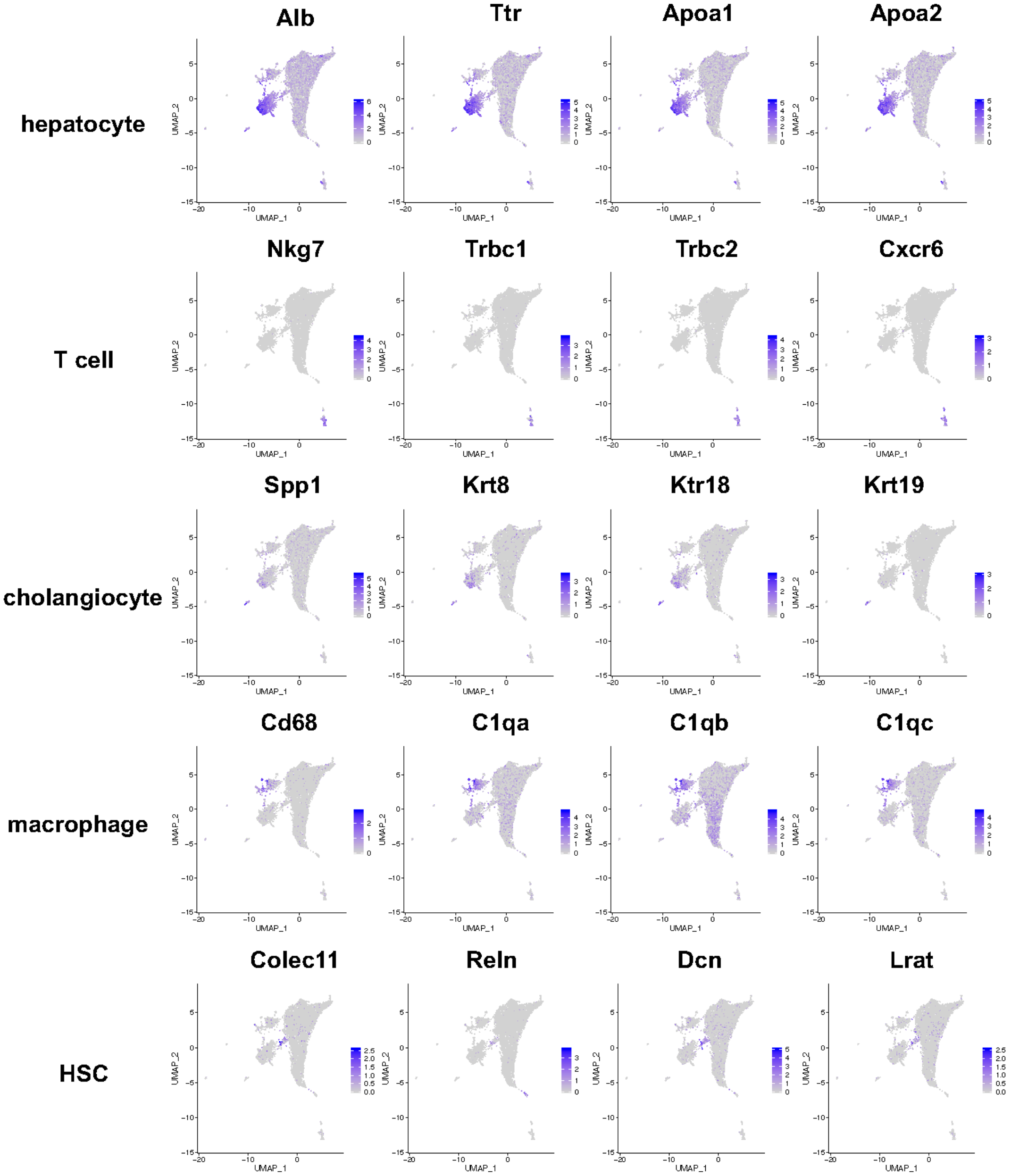


**Supplemental Figure 1 Clusters that expressed non-endothelial cell markers.**
Feature plots showing relative distributions of established marker genes of different liver cell types among the sorted cells.

#### Supplemental Fig 2

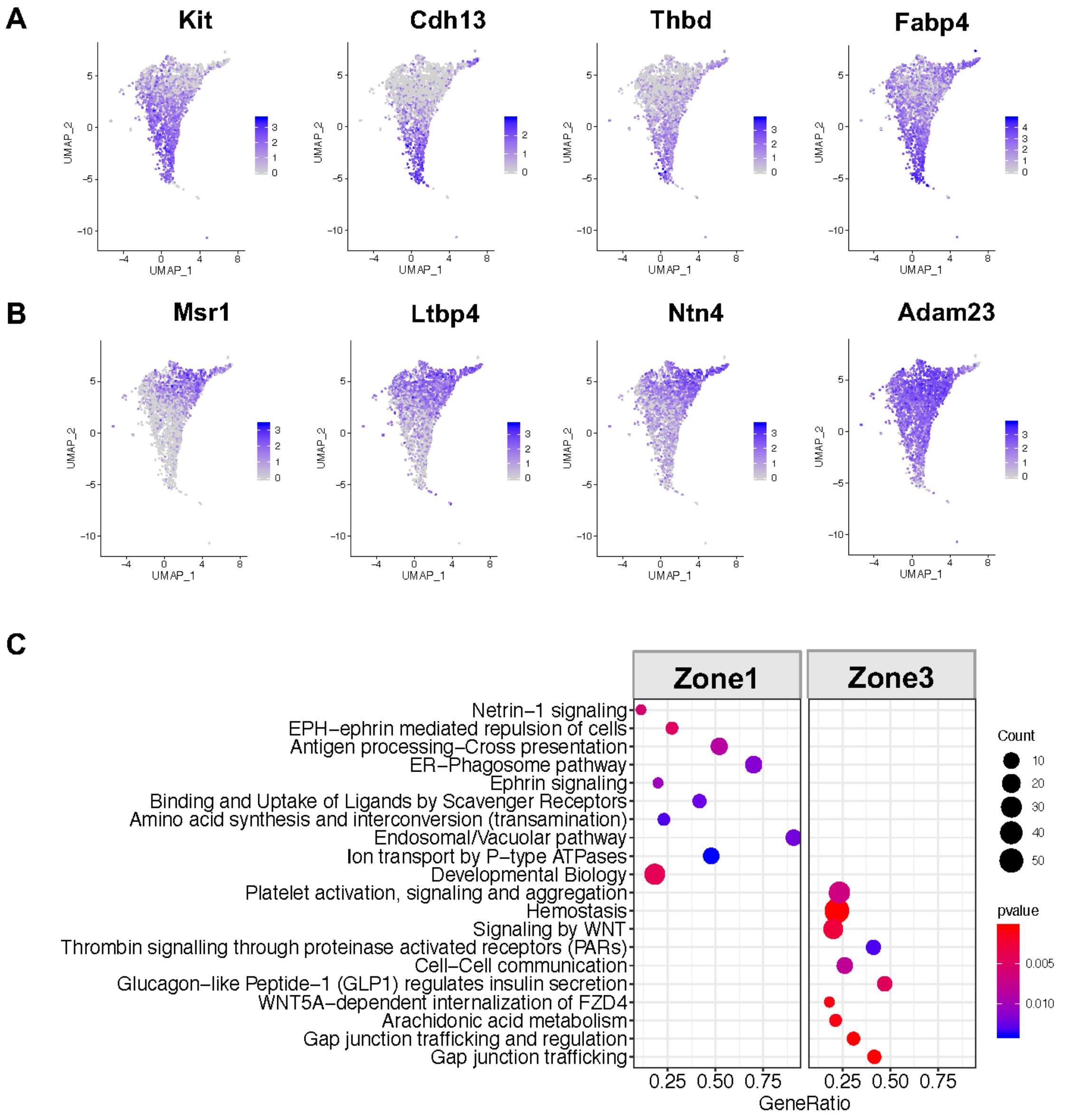


**Supplemental Figure 2 Relative distributions of pericentral and periportal landmark genes among all liver ECs as well as comparison of pathway analysis of periportal and pericentral LSECs.**

**A.** Feature plots showing relative distributions of established pericentral landmark genes among all liver ECs. **B.** Feature plots showing relative distributions of established periportal landmark genes among all liver ECs. **C.** Comparison of signaling pathways enriched in Zone1 LSECs and Zone3 LSECs based on GSEA analysis.

#### Supplemental Fig 3

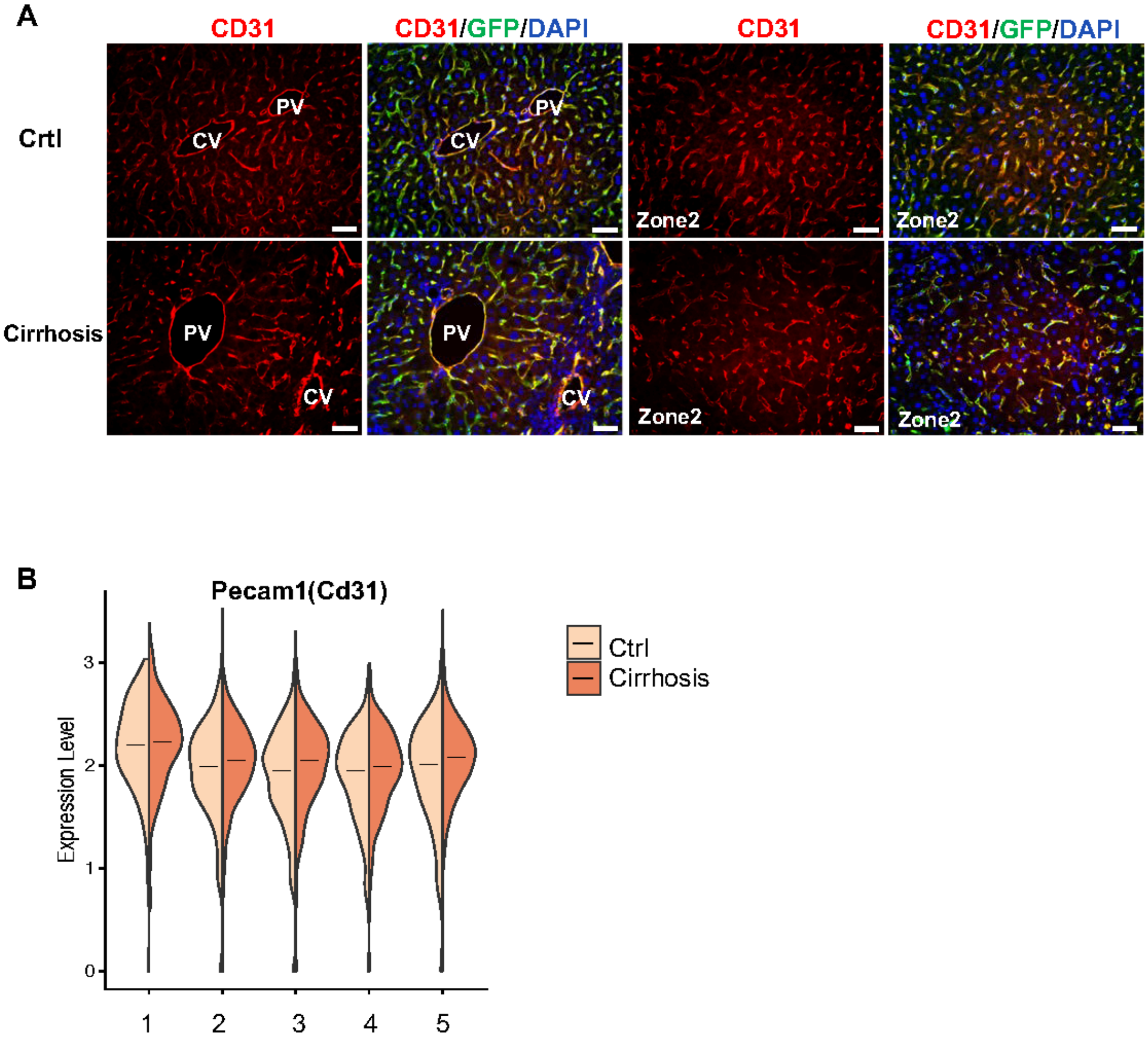


**Supplemental Figure 3 Levels and distributions of CD31 expressing liver ECs are similar between control and cirrhotic mouse livers.**
**A.** Immunofluorescence staining of CD31 (a.k.a., Pecam1) in frozen liver tissue sections from endothelial-GFP reporter mice. Red: CD31, Green: GFP (endothelial cells), Blue: DAPI (nuclei). PV: portal vein, CV: central vein. Scale bar: 40μm Images were taken using a Zeiss fluorescence microscope. **B.** Violin plots showing expression of Pecam1 (Cd31) in liver ECs of Clusters 1-5 in control and cirrhotic mice. Black lines indicate median expression values.

### Supplemental Fig 4

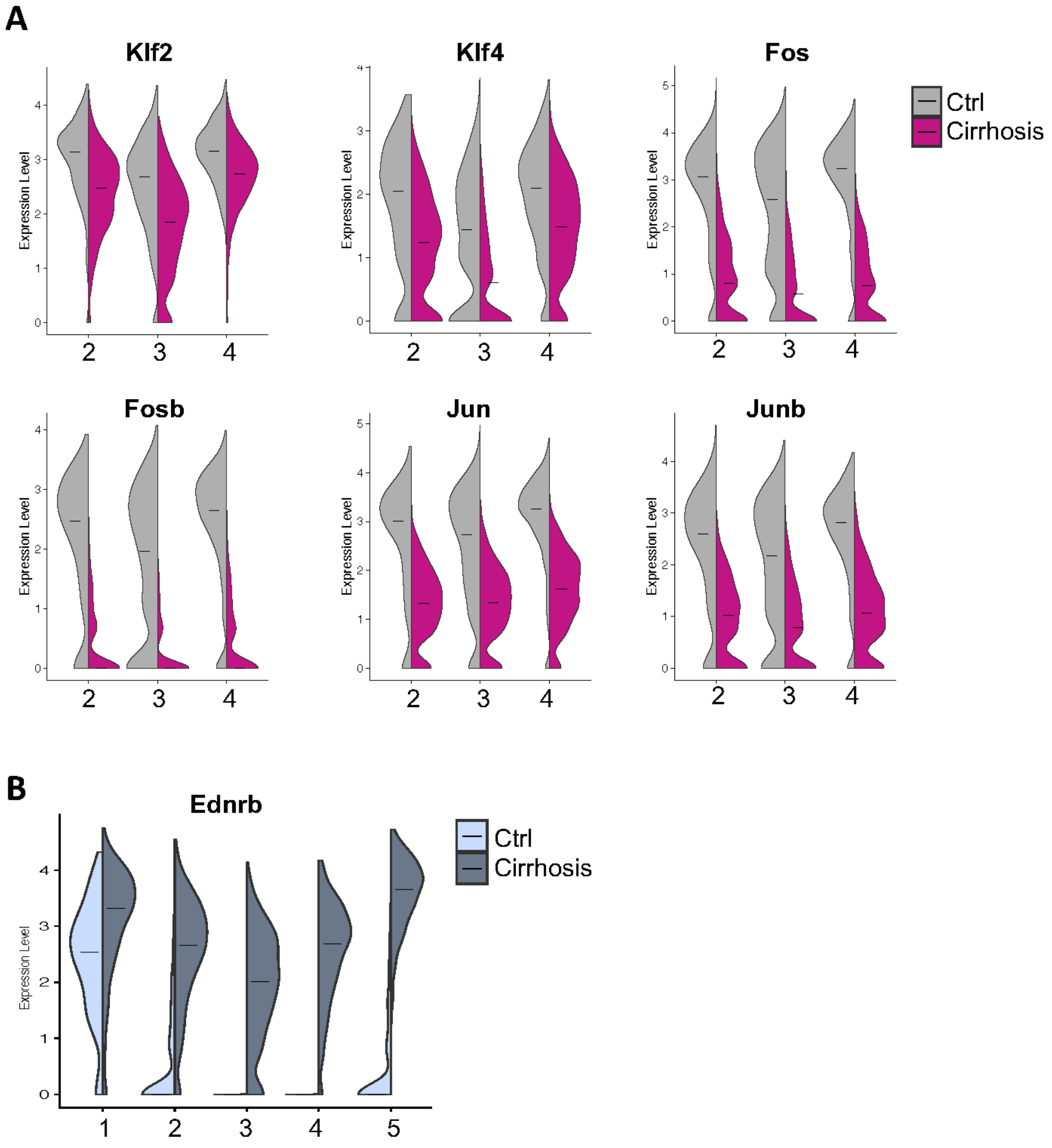


**Supplemental Figure 4 Zonal differences in expression of genes related to EC dysfunction.**
**A.** Violin plots showing expression of transcription factors in LSECs of Clusters 2-4 (corresponding to Zones 1-3) in control and cirrhotic mice. **B.** Violin plots showing expression of Ednrb (endothelin receptor type B) in liver ECs of Clusters 1-5 in control and cirrhotic mice. Black lines indicate median expression values.

#### Supplemental Fig 5

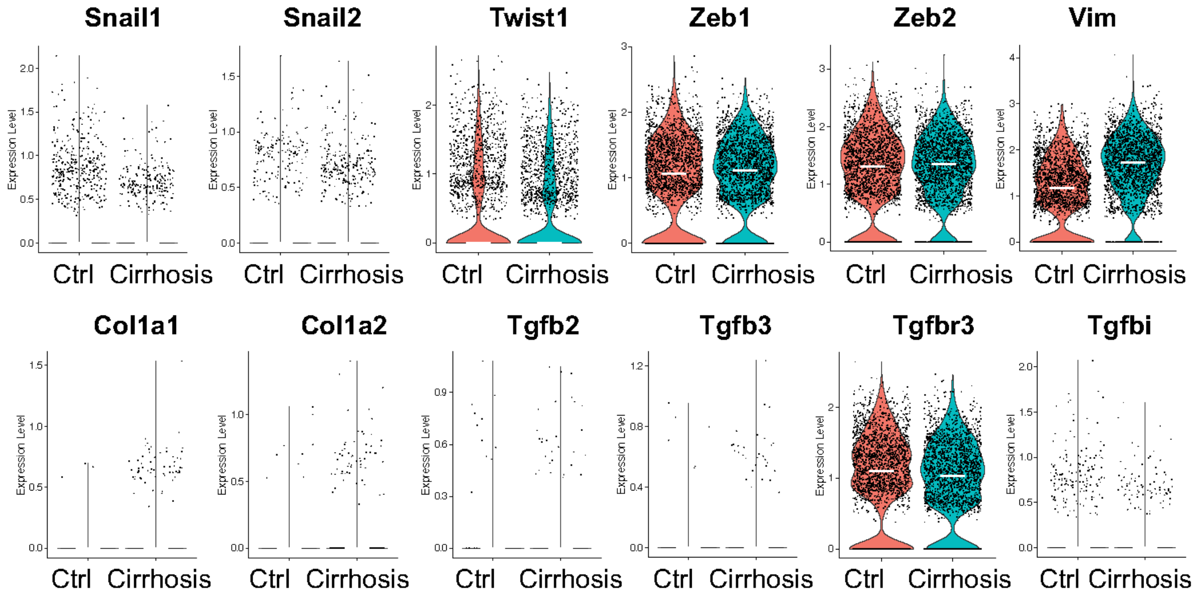


**Supplemental Figure 5 Expression of endothelial-to-mesenchymal transition (EndMT)-associated genes in LSECs (Clusters 2-4) of control and cirrhotic livers.**
Each dot represents a single cell. White lines indicate median expression values.

#### Supplemental Fig 6

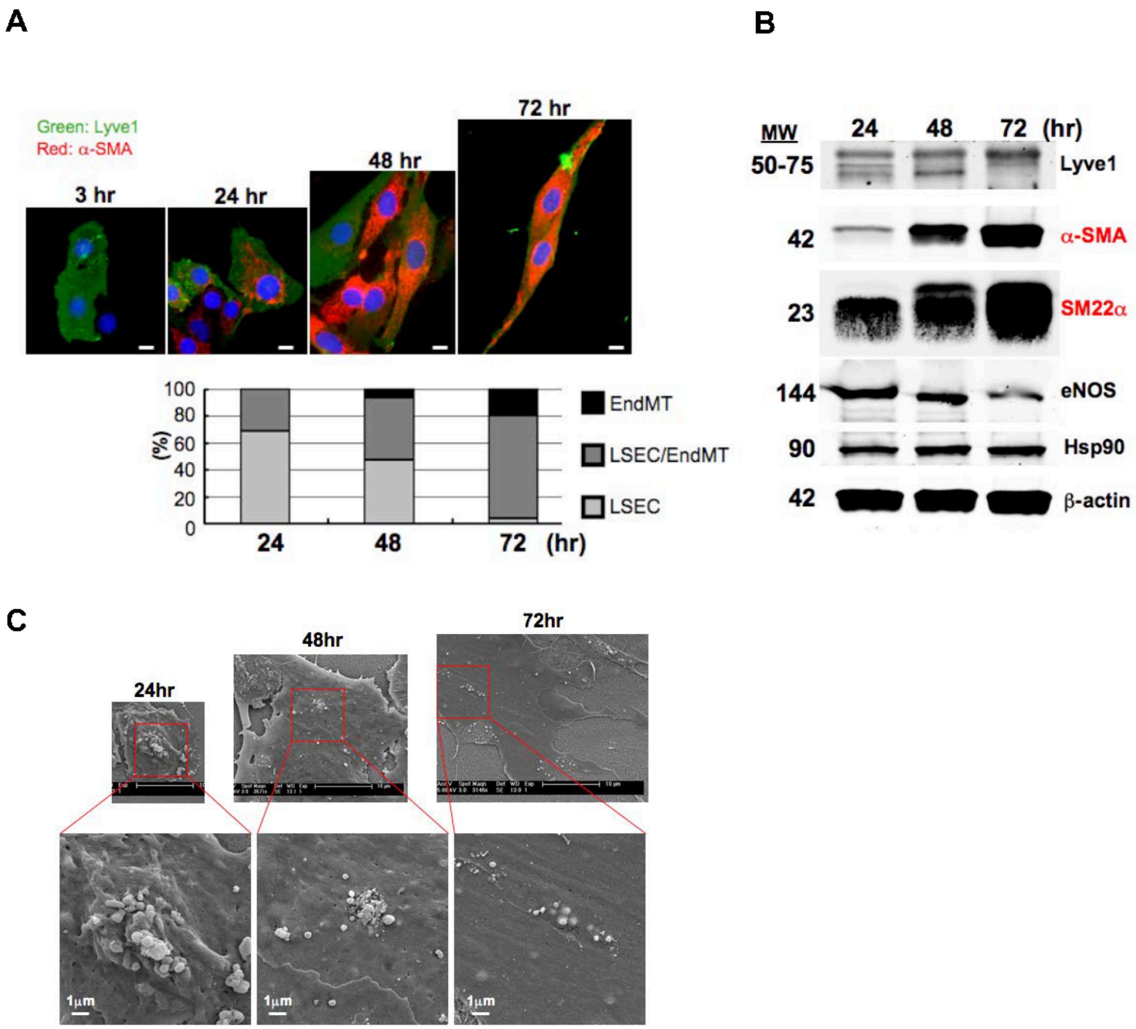


**Supplemental Figure 6 Rat primary LSECs undergo EndMT in a cultured condition in a time dependent manner.**

**A.** Immunofluorescence staining of Lyve1 and alpha-smooth muscle actin (α-SMA) to assess EndMT in rat primary LSECs cultured for 3, 24, 48 and 72 hours on collagen-coated cover glasses. Green: Lyve1 (an LSEC marker), Red: α-SMA (an EndMT marker), Blue: DAPI (nuclei). Scale bar: 5μm. **B.** Western blot analysis of Lyve1 (an LSEC marker), α-SMA and SM22α (EndMT markers) and endothelial nitric oxide synthase (eNOS, an EC marker). Heat shock protein 90 (Hsp90) and β-actin were used as loading controls. **C.** Scanning electron microsopy images of fenestrae in LSECs cultured for 24, 48 and 72 hours. Scale bar: 10μm (upper panel) and 1μm (lower panel).

#### Supplemental Fig 7

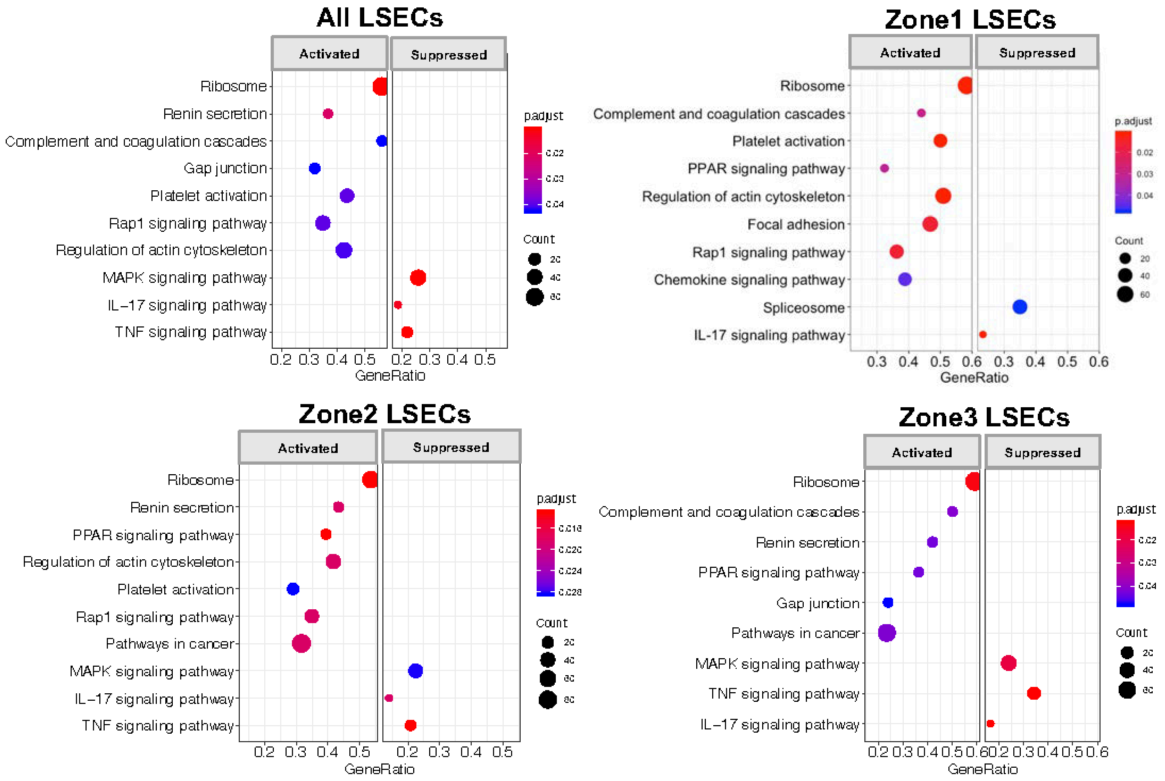


**Supplemental Figure 7 Pathway analysis revealed unique functional changes in zonal LSECs as a result of liver cirrhosis.**
Comparison of signaling pathways by GSEA analysis based on gene expression changes in all the LSECs or within each zone of LSECs in cirrhotic mice compared to control mice.

#### Materials and Methods

##### LSEC and LyEC culture

Primary human liver sinusoidal endothelial cells (LSECs) and lymphatic endothelial cells (LyECs) were purchased from PELOBiotech (Cat#:PB-CH-153-5511, Planegg, Germany) and PromoCell (C 12217, Heidelberg, Germany), respectively. LSECs were seeded on fibronectin-coated cell culture plates and grown in Cellovations® Endothelial Cell Growth Media (PB-MH-100-4099, PELOBiotech) supplemented with 5% FCS and growth factor cocktail according to the manufacturer's instructions. LyECs were seeded on cell culture plates coated with Speed Coating Solution (PB-LU-000-0002-00, PELOBiotech) and cultured in Endothelial Cell Growth Medium MV 2 (C-22221, Promocell) with Growth Medium MV 2 SupplementPack (C-39221, Promocell). All cells were cultured at 37°C in a humidified 5% $CO_2$ atmosphere.

##### Quantitative real-time polymerase chain reaction

Total RNA was extracted from primary human LSECs (PB-CH-153-5511, PELOBiotech) and LyECs (C-12217, Promocell) using Trizol reagent (Invitrogen). Total RNA (1µg) was reverse transcribed into cDNA using Reverse Transcript Reagents kit (04897030001, Roche Molecular Systems, Branchburg, NJ). Quantitative PCR was performed on cDNA with TaqMan Real-time PCR Assays (Thermo Fisher Scientific, Waltham, MA), including human 18S (Hs99999901), human Mmrn1 (Hs01113299), human Prox1 (Hs00896293), human Pdpn (Hs00366766), human Rassf9 (Hs00193763), human Tbx1 (Hs00962558) and human Ahnak2 (Hs00292832). The ABI 7500 real time PCR system (Applied Biosystems, Foster City, CA) was used for amplification.

##### Immunofluorescence

Paraffin sections were de-paraffinized with xylene and rehydrated with graded ethanol. Frozen sections were washed by PBS for 10 min three times. Antigen retrieval was performed using BD solution in a steamer for 20 min. Blocking of nonspecific signal was performed with blocking buffer (5% donkey serum and 0.3% Triton X-100 in PBS) for 1h. Primary antibodies were incubated overnight at 4°C (rabbit anti-α-SMA,1:300, ab124964; rabbit anti-CD34, 1:100, ab81289; rabbit anti-Endrb, 1:100, ab117529, Abcam, Cambridge, MA; rat anti-CD31, 1:100, 550274, BD Pharmingen, San Diego, CA; goat anti-VE-cadherin, 1:200, sc-6458, Santa Cruz Biotechnology). Then, secondary antibodies were incubated for 30 min at room temperature (donkey anti-rabbit Alexa 647, 1:300; donkey anti-rat Alexa 647,1:300; donkey anti-goat Alexa 488, 1:300, Invitrogen). After samples were mounted with FluoroshieldTM containing DAPI (Sigma-Aldrich), their images were taken with a fluorescence microscope (Zeiss Observer Z1, Oberkochen, Germany) or confocal microscope (Leica SP5, Wetzlar, Germany).

##### Isolation of LSEC-enriched fraction (For Supplemental Figure)

Liver non-parenchymal cells (NPCs) were isolated from Sprague Dawley rats as described above for mouse livers[67]. Briefly, after collagenase perfusion and removal of hepatocytes, an NPC fraction was pelleted at 350 xg for 10 min and resuspended in EBM-2 Basal Medium (CC-3156, Lonzs) supplemented with Microvascular Endothelial Cell Growth Medium-2 SingleQuots supplements (CC-4147, Lonza) and 15% fetal bovine serum. The NPC suspension was subjected to a density gradient centrifugation using Percoll (GE Healthcare, Chicago, IL, Lot 10221921) at 900 xg for 20 min at room temperature. An LSEC-enriched fraction was isolated and seeded on collagen-coated coverslips or cell culture dishes. At 3h, 24h, 48h or 72h after cell culture in the endothelial cell medium described above at 37°C in a humidified atmosphere containing 5% $CO_2$, LSECs were used for immunofluorescent staining, Western blot analysis, or scanning electron microscopy.

**Immunocytochemistry of LSECs (For Supplemental Figure)**
LSECs seeded on cover glasses were fixed with 4% paraformaldehyde (PFA) for 20 min, permeabilized with 0.1% Triton X-100 for 15 min, blocked with 5% donkey serum for 60 min, and incubated with primary antibodies in a humidified chamber overnight. Rabbit anti-LYVE1 (1:100, ab14917, Abcam) and mouse anti-α-SMA (1:300, M0851, Agilent Dako, Santa Clara, CA) were used as the primary antibodies.

**Western blot (For Supplemental Figure)**
Proteins were extracted from LSECs using a lysis buffer containing 50mmol/L Tris-HCl, 0.1mmol/L EGTA, 0.1mmol/L EDTA, 0.1% SDS, 0.1% deoxycholic acid, 1% (vol/vol) Nonidet P-40, 5mmol/L sodium fluoride, 1mmol/L sodium pyrophosphate, 1mmol/L activated sodium vanadate, 0.32% protease inhibitor cocktail (Roche Diagnostics, Mannheim, Germany), and 0.027% Pefabloc (Roche Diagnostics). Protein concentrations were measured using a modified Lowry assay method with DC protein assay reagents (Bio-Rad Laboratories, Hercules, CA). 20 μg of protein was loaded and separated by SDS-PAGE. Proteins transferred to 0.2μm nitrocellulose membranes (Bio-Rad Laboratories) were analyzed by immunoblotting with primary antibodies including rabbit anti-α-SMA (1:3000, ab124964, Abcam), rabbit anti-SM22α (1:3000, ab14106, Abcam), mouse anti-eNOS (1:1000, 610297, BD Biosciences), mouse anti-HSP90 (1:1000, 610419, BD Biosciences) and mouse anti-β-actin (1:3000, A1978, Sigma-Aldrich). After washing with Tris buffered saline containing 0.1% Tween-20 (TBS/T), membranes were incubated with fluorophore-conjugated secondary antibodies (LI-COR Biotechnology, Lincoln, NE) having 680nm or 800nm emission. Proteins were visualized and quantified using the Odyssey Infrared Imaging System (LI-COR Biotechnology). Heat shock protein 90 (Hsp90) and β-actin were used as loading controls.

**Scanning electron microscopy (For Supplemental Figure)**
LSECs were seeded on collagen-coated cover glasses in twelve-well tissue culture plates. After 24h, 48h and 72h, LSECs were fixed with 2.5% glutaraldehyde in 0.1M cacodylate buffer with pH7.4 at room temperature for 30 min and then moved to 4°C for 1h. After washing with PBS, LSECs were treated with 1% tannic acid in 0.15M cacodylate buffer for 1 h, then fixed with 1% osmium tetroxide in 0.1M cacodylate buffer for 30 min, dehydrated with graded alcohols, dried with hexamethyldisilazane, and examined using a scanning electron microscope (Hitachi SU-70, Tokyo, Japan).
